# Different approaches to processing environmental DNA samples in turbid waters have distinct effects for fish, bacterial and archaea communities

**DOI:** 10.1101/2022.06.17.495388

**Authors:** Rachel Turba, Glory H. Thai, David K. Jacobs

**Affiliations:** Department of Ecology and Evolutionary Biology, University of California, Los Angeles, California 90095 USA

**Keywords:** metabarcoding, turbidity, method comparison, fish

## Abstract

Coastal lagoons are an important habitat for endemic and threatened species in California that have suffered impacts from urbanization and increased drought. Environmental DNA has been promoted as a way to aid in the monitoring of biological communities, but much remains to be understood on the biases introduced by different protocols meant to overcome challenges presented by unique systems under study. Turbid water is one methodologic challenge to eDNA recovery in these systems as it quickly clogs filters, preventing timely processing of samples. We investigated biases in community composition produced by two solutions to overcome slow filtration due to turbidity: freezing of water prior to filtration (for storage purposes and long-term processing), and use of sediment (as opposed to water samples). Bias assessments of community composition in downstream eDNA analysis was conducted for two sets of primers, 12S (fish) and 16S (bacteria and archaea). Our results show that freezing water prior to filtration had different effects on community composition for each primer, especially for the 16S, when using a filter of larger pore size (3 μm). Nevertheless, pre-freezing water samples can still be a viable alternative for storage and processing of turbid water samples when focusing on fish communities (12S). The use of sediment samples as an alternative to processing water samples should be done with caution, and at minimum the number of biological replicates and/or volume sampled should be increased.

## Introduction

Coastal lagoons in California are the numerically dominant form of coastal wetland (Jacobs et al., 2011; Stein et al., 2014) and are important in many other Mediterranean climates and subtropical environments. These lagoons are characterized by seasonal and episodic breaching (opening of the lagoon to the sea, usually by stream flow) and closure (isolation of the lagoon by a high sandbar), which provide a suite of ecological services: from groundwater infiltration to support of unique biodiversity (Ballard et al., n.d.). This system serves as important habitat and nursery for endemic and endangered fishes and amphibians, such as the steelhead (*Oncorhynchus mykiss*), red-legged frog (*Rana aurora draytonii*), and the tidewater goby (*Eucyclogobius newberryi*) (Earl et al., 2010; Shaffer et al., 2004; Swift et al., 1993, 2016). Thus, California lagoons are spatially and temporally variable systems with unique biodiversity and biodiversity assessment challenges.

Coastal lagoons have been drastically reduced in numbers along the California coastline, driven mostly by the impact of coastal land use for transport structures, agriculture, and development. These are further exacerbated by ongoing changes in the hydrological cycles due to climate change (SCWRP, 2018). While these sites are critical for endangered species conservation, they are also subject to frequent invasion and their response to environmental variation is poorly documented. However, monitoring of this habitat can be limited by a variety of issues, ranging from limited human power and access to challenges driven by the natural complexity and dynamism of these lagoons.

The use of environmental DNA (eDNA) has been advocated as an alternative for monitoring communities and target species (Thomsen & Willerslev, 2015), and can overcome and complement certain field limitations from traditional methods (e.g. seining, trapping). On-site collection can be relatively fast, and therefore allow field workers to cover more ground. It can also recover the DNA signal of species that are rare, cryptic and/or hard to capture by traditional methods, and being non-intrusive, it offers an alternative when working with endangered species for which permits are necessary (Deiner et al., 2017; Dejean et al., 2012; Sard et al., 2019). In addition, metabarcoding approaches allow the investigation of multiple species from a single collection (Taberlet, Coissac, et al., 2012).

Nevertheless, it is important to recognize that this approach also brings its own limitations and biases (van der Loos & Nijland, 2021). In some circumstances, eDNA sampling can be more expensive than traditional, more established methods (Smart et al., 2016). Since there are no voucher specimens from collections, contamination is a major issue that needs to be addressed early on, following best practices in the field (Goldberg et al., 2016). The lack of voucher specimens also leads to an overdependence on the use of barcodes and genetic databases for taxonomic identification, which introduces another set of biases, from misidentification to lack of species representation (Taberlet, Coissac, et al., 2012). Other challenges arise from the non-universality of sampling methods and downstream processing, with the probability of detection varying depending on the species and their density, as well as the type of environment, which affects rates of DNA degradation (Deiner et al., 2015; Nagarajan et al., 2022; Rees et al., 2014; Williams et al., 2017).

Coastal lagoons can vary drastically in their environmental properties. One major challenge is the high and variable turbidity of the water. High turbidity often occurs during high stream flow, or when lagoons are closed to the ocean by a sandbar leading to an accumulation of organic and inorganic matter. In this scenario, filtering turbid water on-site is problematic.

However, filtration is widely used for handling water samples (Laramie et al., 2015; Tsuji et al., 2019). Typically, set volumes of water are run through a small pore size filter to concentrate DNA before extractions. However, high concentration of fine sediment or organic matter in water quickly obstructs these filters, making the filtration process time-consuming (although it could actually aid recovery by binding DNA to suspended particles: Kumar et al., 2022; Liang & Keeley, 2013; Torti et al., 2015).

To overcome this issue, some stakeholders have relied on a tiered filtration step (prefiltration) to reduce particles and avoid clogging filters (Tsuji et al., 2019), but this approach increases costs, labor and opportunities for potential contamination (Li et al., 2018; Majaneva et al., 2018; Robson et al., 2016). The use of filters with bigger pore sizes (up to 20 µm) has been previously tested and in cases of turbid waters is generally preferred, but requires filtering larger volumes of water to capture the same amount of DNA recovered in smaller pore size filters (Robson et al., 2016; Turner et al., 2014).

Freezing water for storage purposes prior to filtration can mitigate the issue of slow filtration in the field and allow it to be done in batches in the laboratory at a later time, but this type of sample storage could also introduce bias on eDNA capture and community composition (Kwambana et al., 2011; Sekar et al., 2009). Freezing can disrupt cells and extrude their DNA in the environment, which would then make it easier for it to pass through the filter pores and be lost, an issue that has been demonstrated in certain cases (e.g. Suomalainen et al., 2006).

Increasing the pore size of filters to speed the filtration process could worsen this problem by letting DNA in solution flow through the pores more easily.

When dealing with turbid waters, some stakeholders have opted to use the centrifugation approach (e.g. Williams et al., 2017). Extracellular DNA (i.e. DNA not contained within a cell wall) can be bound to particles (Torti et al., 2015) and consequently be captured and detected following centrifugation of particles into pellets. However, the amount of water used is limited by centrifuge size, usually around 15-30 mL per replicate (Doi et al., 2017; Ficetola et al., 2008), which might limit recovery of diluted DNA (Deiner et al., 2015).

Processing sediment samples may be preferable to processing highly turbid water samples. However, it is important to understand how DNA recovery from these different media compare to one another. Turner et al. (2015) and Perkins et al. (2014) have shown that sediment can have a higher concentration of fish eDNA and some bacteria, respectively. This may relate to the organic-particle binding and sinking properties, and longer persistence of DNA in sediment compared to water samples. However, as is the case with water samples, there is no consensus on the rate of degradation of eDNA in soil and sediment (Dell’Anno & Corinaldesi, 2004; Levy-Booth et al., 2007; Torti et al., 2015), and this will depend on multiple local biotic and abiotic factors. In addition, biological communities can naturally differ between water column and sediments, even when we expect some level of overlap due to both DNA sinking and suspension.

Previous works have compared different approaches to processing eDNA, including assessment of filtration and storage methods (Hinlo et al., 2017; Takahara et al., 2015), comparisons between water and sediment eDNA recovery (Sales et al., 2019; Turner et al., 2015), as well as some work on turbid waters (Kumar et al., 2022; Robson et al., 2016; Williams et al., 2017). But results have been contradictory, or limited to looking at just DNA concentration, or at a single targeted species.

The goal of the present study is to compare how freezing water prior to filtration and using water versus sediment samples induce and/or exacerbate biases in taxa detection for a set of universal primers targeting different biological communities–12S (fish) and 16S (bacteria and archaea)—in coastal lagoons. By understanding the biases introduced when processing environmental samples, we will be able to inform decisions regarding experimental design for monitoring such a dynamic and challenging habitat, which has invaluable importance for the maintenance of ecosystem services for both wild and urban populations. We expect these results will be of interest relative to eDNA sampling in other aquatic systems as well, such as rivers, streams, and ponds, especially those with turbid waters.

## Material and Methods

### Site - Topanga Lagoon

To determine the variability of species detection for each protocol, water and sediment samples were collected from a south-facing coastal lagoon in southern California, located in Malibu, a stretch of coast that runs from Santa Monica to Point Mugu. This lagoon is part of the Topanga State Park and is currently undergoing plannings for restoration. It is the only lagoon on this stretch of coast that still harbors a stable population of tidewater goby (*E. newberryi*), a federally endangered species, and is relatively less impacted than other lagoons in the same region. The endangered southern steelhead trout (*O. mykiss*) is also found in this system during anadromy when the lagoon is breached. Due to the presence of these species, Topanga lagoon has been periodically surveyed by the Jacobs’ lab members and collaborators such as researchers at the Resource Conservation District of The Santa Monica Mountains (RCDSMM), and therefore its macrobiota is regularly studied, especially the fish fauna. The lagoon was sampled on September 6th, 2018, at the end of the Summer season, and as is typical of this time of the year, the weather was dry with no record of precipitation since June (WeatherSpark.com, n.d.). The lagoon was closed to the ocean by a sandbar and the water was murky (Fig. 1), which in the author’s experience, such turbidity slowed filtration and easily clogged 0.45 μm cellulose nitrate filters.

**Figure 1:**
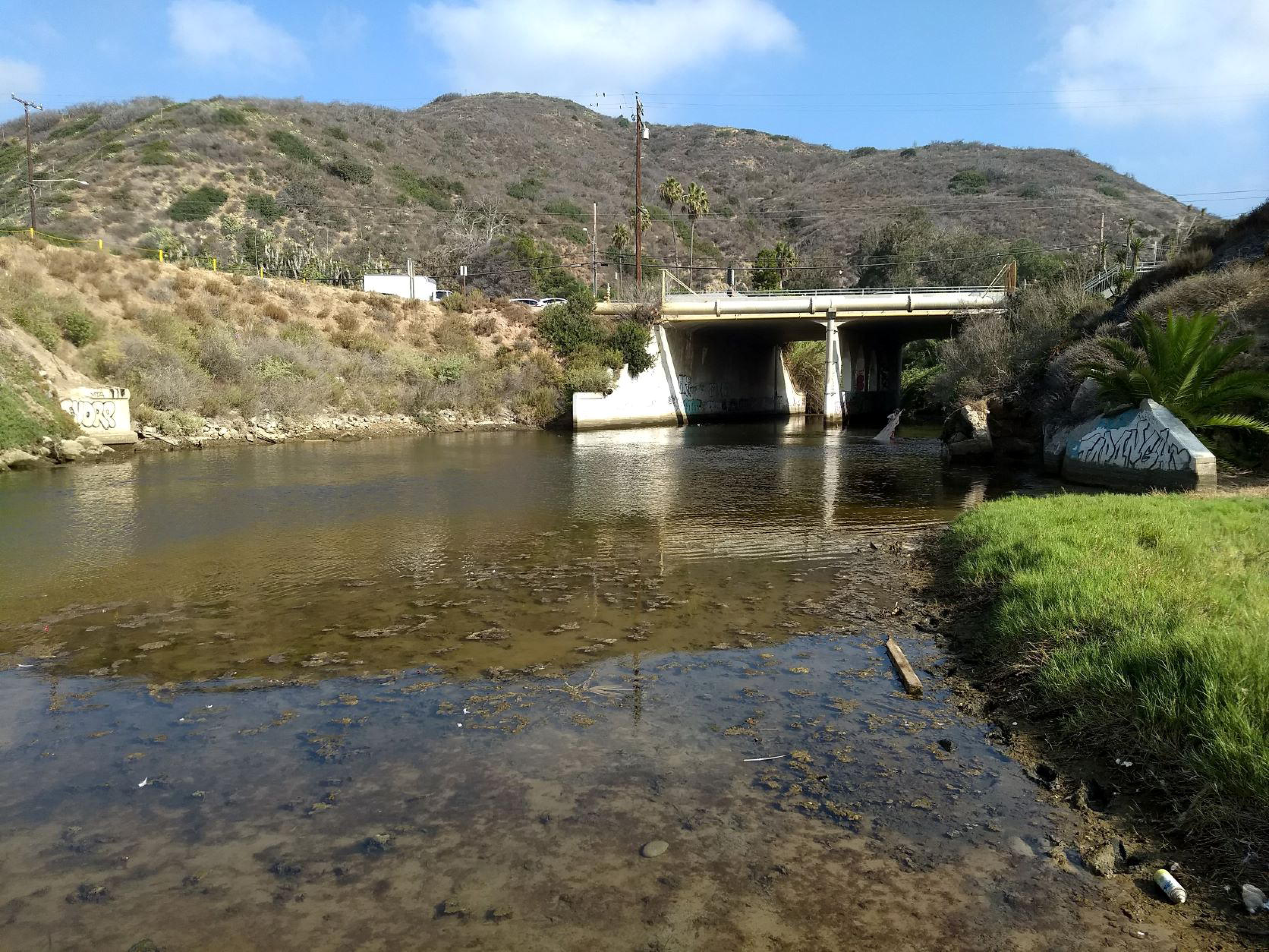
Photo of Topanga lagoon taken on August 22nd, 2018, a few weeks after collection. There was no record of precipitation for the previous three months and the lagoon was closed to the ocean by a sandbar. There was also no sign of recent waves topping over the sandbar and reaching the lagoon.

### Protocols and samples

A sterilized water jug was used to collect a single water sample in the lagoon, at a mid-point between the mouth margin and the road bridge (Fig. 1). The sample was then placed on ice and brought to the laboratory (∼1 hr car ride). This method of “grab-and-hold” has proven to be similarly effective as on-site filtration in a previous study (Pilliod et al., 2013). Once in the laboratory, the total volume was divided for two separate protocols: centrifugation; and filtration.

The centrifugation protocol followed Doi et al. (2017) using five replicates of 50 mL tubes (with 27 mL water samples each). Besides extracting the pellet, we also included a filtration step of the supernatant using a 0.45 µm filter. For the filtration protocol, water was separated into ten 500 mL bottles (Fig. 2) and these were used in two separate protocols. Half (i.e. five bottles) were frozen in the -20°C for three days before thawing for filtration (hereafter referred to ‘pre-freezing (PF) protocol’), and half was not frozen and immediately filtered (hereafter referred to ‘no freezing (NF) protocol’). Filtration was done in two sequential steps for both treatments (pre-and no freezing) using an adapted vacuum pump in the pre-PCR room of the laboratory (Fig. S1). First, the water sample was filtered using a 3 µm filter, then the filtrate was passed through a 0.45 µm filter. All filters used in this work were cellulose nitrate. Here, however, we will focus only on the results from the first filtration step of the water filtration protocol (3 µm filters). More details on that are further explained in the supplemental material.

**Figure 2:**
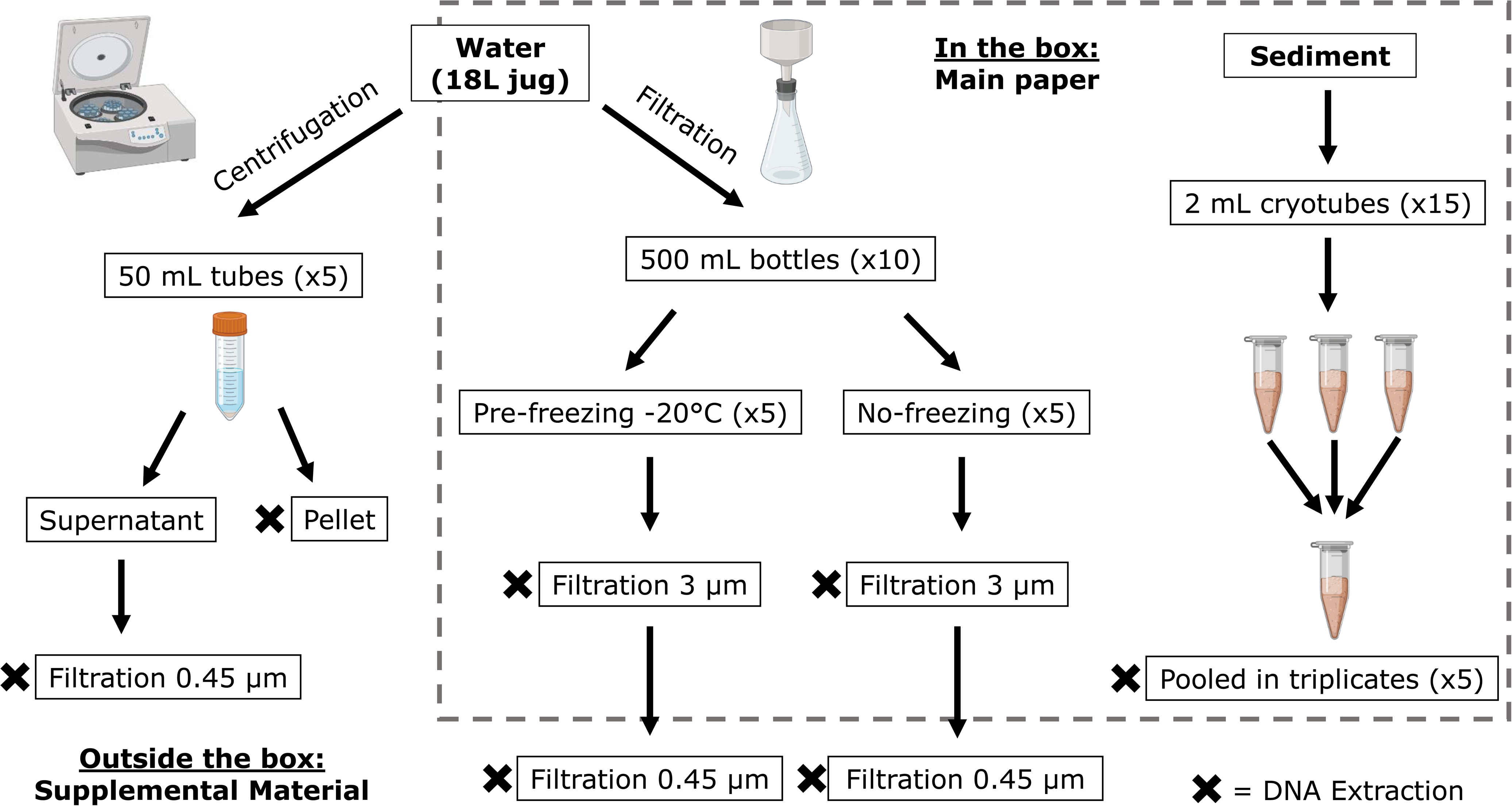
Flowchart of the study design. Water was taken as a single sample in a 18 L water jug from the middle of the lagoon (Fig. 1). In the laboratory, the water was split in batches to follow either the centrifugation or the filtration protocol. Sediment samples were taken in 2 μL cryotubes at the same location where water was taken. Subsamples of these were then pooled in triplicates prior to extraction (total 0.25-0.3 g). The dotted box indicates which part of the study design is addressed in the main paper, and which is addressed in the supplemental material. The black ‘X’ indicates the stage at which DNA extraction was performed and sequenced. Figures were obtained from Biorender.

Surficial sediment (within the first 5 cm) was also collected at the same location where water was sampled (hereafter referred to ‘sediment (Sed) protocol’). We used five collection kits, and each kit consisted of three 2 mL tubes (15 tubes total). Subsamples of these were pooled in triplicates prior to DNA extraction following instructions as defined by the CALeDNA program (https://ucedna.com/methods-for-researchers). These were also kept on ice during field work and stored in a -80°C freezer upon arrival at the laboratory until DNA extractions.

In order to test if freezing water is a viable process to store and manage water samples, we compared the results of pre-freezing water prior to filtration with the no freezing protocol. Results from the sediment protocol were compared against both filtration protocols: (pre-freezing versus no freezing) to test if sediments can be used as an alternative to water samples.

### DNA Extraction

DNA from sediments and filters were extracted following the PowerSoil extraction protocol. Filters were chopped into thin strips before being added to the bead tubes, and sediment triplicates were pooled in small batches to reach 0.25-0.3 g before processing. We used the soil extraction kit on the filters as well to reduce potential PCR inhibition caused by the water turbidity (Kumar et al., 2022), but also to limit the number of variables in the research design by adding another extraction protocol.

### Contamination best practices

Care was taken to avoid contamination both in the field and the lab. Before collection, bottles and the water jug were cleaned and bleached and then handled with clean gloves on site. Extractions and PCR were done in a separate pre-PCR room. Utensils and bench top were cleaned with 10% bleach, followed by 70% ethanol. Forceps and scissors for handling filters were seared and cleaned with bleach and ethanol after dealing with each sample. PCR reagents were prepared in a clean, PCR-free, positive pressure hood. Sediment samples were collected with new 2 mL cryotubes and following field protocol as recommended by the CALeDNA program. Blanks were made for the field collection, laboratory filtration and PCR (5 blanks in total) and included in the library for sequencing.

### Sequencing

Library preparation followed CALeDNA protocols (https://ucedna.com/methods-for-researchers). Metabarcode libraries were generated for bacteria and archaea (16S rRNA), fish (12S rRNA) and metazoans (CO1). Sequences for each primer can be found at Table 1. All libraries consisted of triplicate PCR reactions. PCR products were visualized using gel electrophoresis, and for each barcode, PCR triplicates were pooled by sample. After bead cleaning, all markers were pooled by sample and tagged for sequencing (single indexing). Libraries were pooled and run on a MiSeq SBS Sequencing v3 in a pair-end 2×300 bp format [Technology Center for Genomics & Bioinformatics (TCGB), UCLA] with a target sequencing depth of 25,000 reads/sample/metabarcode. Two sequencing runs were conducted, but the CO1 primer was still below the sequencing depth threshold and therefore its results will not be discussed here (see Figs. S2-3). For each run, our library was pooled with different samples from different collaborators to maximize efficiency of the sequencing run.

**Table 1:**
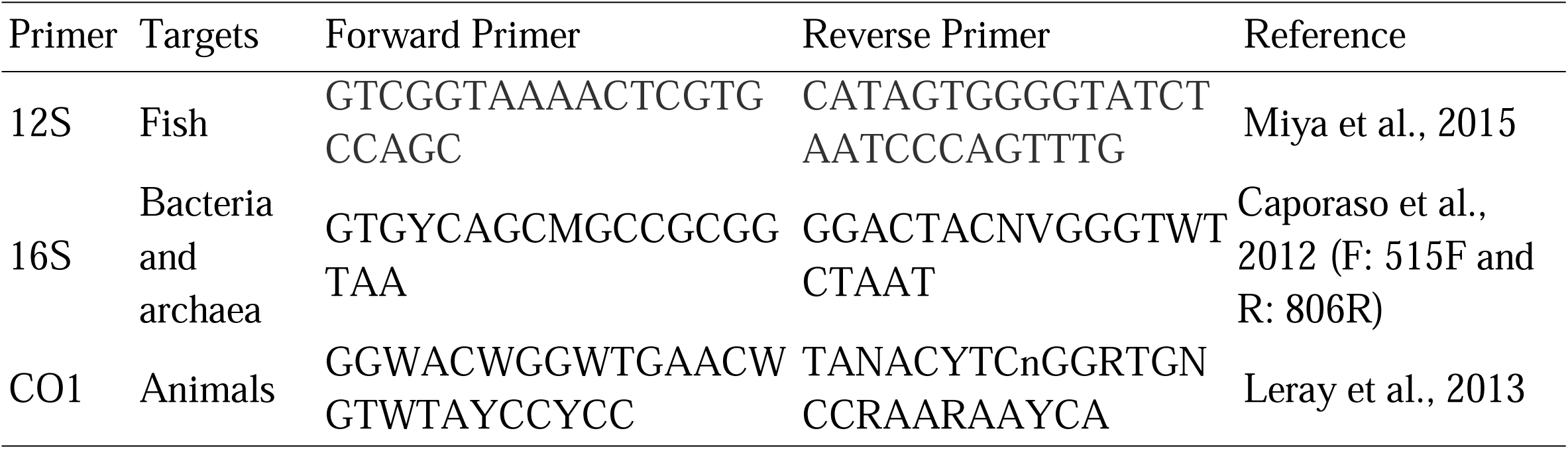
Detailed information of the primers used.

### Bioinformatics and data pre-processing

Sequence data was bioinformatically processed in Hoffman2, the High Performance Computing cluster at UC Los Angeles, using the Anacapa Toolkit (Curd, Gomer, et al., 2018) with default settings. Briefly, reads are demultiplexed and trimmed for adapters (cutadapt, Martin, 2013) and low-quality reads (FastX Toolkit, *FASTX-Toolkit*, n.d.). Dada2 (Callahan et al., 2016) is used to denoise, dereplicate, merge and remove chimeras, and the resulting clean Amplicon Sequence Variants (ASVs) have their taxonomy assigned using Bowtie2 (Langmead & Salzberg, 2012), matched to a custom reference library (CRUX, Curd, Kandlikar, et al., 2018). Confidence levels are determined by the BLCA algorithm (Gao et al., 2017) to generate a table of best taxonomic hits, from super-kingdom to species level. The pipeline was designed to process not only paired, but also unmerged and unpaired reads.

Taxonomic tables with a bootstrap confidence cutoff score of 0.6 were used for downstream analyses. Except when noted, all bioinformatic analyses mentioned beyond this point were performed using R v.3.6.2 (R Core Team, 2018) in RStudio v.1.2.1335 (RStudio Team, 2020). Decontamination was done separately for each primer set and each run (since the dataset was pooled with different combinations of samples for sequencing). We used the package metabaR (Zinger et al., 2020) to lower tag-jumping and remove contaminants through detection of ASVs whose relative abundance is highest in negative controls. We also ran a modification of the gruinard pipeline (https://github.com/zjgold/gruinard_decon), including only steps 4 (site occupancy modeling) and 5 (dissimilarity between replicates), since previous steps were redundant with the metabaR decontamination steps. Lastly, taxa classified as “Not_found”, “Unclassified”, “*Canis lupus*”, “*Bos taurus*”, and “*Homo sapiens*” were removed from the final tables before being merged and used in downstream analyses.

### Diversity analysis

We used the laboratory’s own sampling record and the Global Biodiversity Information Facility database (Gbif.Org, 2022) to manually check the 12S primer final taxonomic table. The number of species captured by each protocol was visualized using Venn Diagrams (package VennDiagram, Chen, 2018). Species rarefaction curves were made for each metabarcode to inspect the level of species saturation in each protocol replicate. The slope of each curve was calculated using the rareslope function in the vegan package (Oksanen et al., 2019), and the confidence interval was calculated using pairwiseCI (Schaarschmidt & Gerhard, 2019) with confidence level at 95%. Rarefaction curves were plotted using the ggrare function from the ranacapa package (using step = 5).

### Differential abundance

The raw dataset was analyzed using DESeq2 and ALDEx2 to look at differential abundance between protocols (Fernandes et al., 2013; Love et al., 2014), and the output of both analyses were compared to look for consistency in results. While DESeq2 uses a series of negative-binomial generalized linear models to count data and estimate the log2-fold change in abundance, ALDEx2 (ANOVA-like Differential Expression) accounts for community composition when calculating differential expression by performing a scale invariant transformation on read counts, which are modeled as distributions of posterior probabilities sampled from a Dirichlet distribution. This method has been found to produce more consistent and reproducible results (Nearing et al., 2022).

For DESeq2, the default testing framework was used (test = “Wald”, fitType = “parametric”), which includes the Benjamini-Hochberg multiple inference correction. The sfType option was defined as poscounts since this estimator is able to handle zeros. The log2 fold change of each pairwise comparison for which there were significant differences in abundances was plotted. For ALDEx2, we performed a two sample T-test for each pairwise comparison: between the different filtration protocols (pre-versus no freezing), and between each filtration protocol and sediment protocol (i.e. PF vs. Sed; and NF vs. Sed). We used 500 Monte Carlo samples (mc.samples = 500) for estimation of the posterior distribution. We plotted both the between-(M) versus within-condition fold change (W) (MW) and Bland-Altman (BA) plots using the aldex.plot function.

### Beta diversity

For the beta diversity analysis, samples were standardized by using either the eDNA index (Kelly et al., 2019) or by rarefying them as a way to equalize sequencing effort and minimize stochasticity and bias. For the eDNA index, we followed the Wisconsin double standardization method in the vegan package. The custom_rarefaction function in the R package ranacapa (Kandlikar, 2020) was used to rarefy the dataset with 10 replicates.

For the 12S primer, samples were rarefied to 20 000 reads. Three sediment replicates were excluded due to very low read numbers (<100). For the 16S, samples were rarefied to 15 000 and one sediment replicate that had ∼5000 reads was excluded. The number of reads per taxa for each protocol replicate was plotted using the phyloseq package (McMurdie & Holmes, 2013), for both the raw and rarefied dataset.

The rarefied dataset followed a Constrained Analysis of Principal Coordinates (CAP) using the capscale function in vegan and Bray-Curtis distance. This ordination method, which can be used with non-Euclidean dissimilarity indices, explains the ordination of assemblage composition based on species abundances. The difference in community composition for each treatment was then analyzed using a PERMANOVA and Bray-Curtis dissimilarity, followed by a pairwise PERMANOVA comparison (all with the vegan package). P-values were adjusted using the FDR (False Discovery Rate) approach.

## Results

### Sequencing

The first run generated a total of 6 407 371 reads: 3 817 216 reads for the 12S primer, 2 393 627 for 16S, and 196 528 for CO1. In the second run there were a total of 9 088 496 reads: 6 685 673 reads for the 12S metabarcode, 1 904 283 reads for 16S and 498 540 for the CO1. For the 12S and 16S primers, we were able to reach our threshold of 25 000 reads/sample in most cases, while that was not the case for all except one sample of the CO1 primer. Because of this limitation on the number of reads/sample, the CO1 metabarcode will not be discussed further in the main paper (but check the supplemental material for more details, Turba et al., 2023).

### Bioinformatics and data pre-processing

The number of reads per sample after decontamination and combining both runs is illustrated in Figure S3. We manually checked the final taxonomic tables of each separate run for the 12S primer to look for signs of contamination and evaluate how well the bioinformatic decontamination steps worked (metabaR and gruinard). The taxonomic tables for the 12S primer have substantially less species than the 16S, and the local fish fauna is relatively well known, making the process more tractable.

For the run that was pooled with samples from Palmyra Atoll, the output still retained some tropical reef and pelagic fish and elasmobranch species that are not found in coastal lagoons in California. We can expect that tag-jumping contamination is also present in the other sequencing runs and primers as well. Interestingly, eight out of 28 of those tropical species (ca. 28%) were found exclusively on the sediment protocol and not the filtration protocols (e.g. *Acanthurus achilles, Scarus altipinnis, Lutjanus russellii*).

Barplots for both the raw and rarefied dataset (Figs. S3-4, respectively) show that sediment replicates had greater variability amongst themselves, both in number of reads and community composition, compared to the replicates of either filtration protocols. Water replicates were more consistent within and between protocols, and had an overall higher number of reads than the sediment replicates.

### Diversity

After the decontamination steps (metabaR and gruinard) and removing specific, uninformative ASVs (as listed above), the total number of species assigned to 12S was 39, distributed in 20 orders and 22 families. Of these 39 species, only four had been previously recorded for the site (Table S1). For 16S, the total number of taxa assigned to species was 2 625, distributed in 45 phyla and 335 families.

We also noticed some dubious taxonomic assignments. For example, for the 12S primer, we had one hit for *Fundulus diaphanus*, which is a species of killifish native to the northeast of North America. However, the californian species *F. parvipinnis* has been previously documented in Topanga by lab members sampling at the site. Similarly, there were two hits for *Phoxinus phoxinus*, which has a European distribution with a closely related North American counterpart, *P. eos*, although this species has not been identified in collections from Topanga lagoon. Another dubious identification occurred for two species of *Odontesthes*, *O. incisa* and *O. smitti*, which were among the most abundant hits in our dataset but are native to the southwest Atlantic. These two species, however, are South American relatives of topsmelt (*Atherinops affinis*), commonly found in coastal lagoons and estuaries in California (Table S1).

The Venn Diagram (Fig. 3) shows that even though the sediment protocol had lower numbers of reads overall (Figs. S2-3), they had the highest number of species recovered (12S primer: N=27, 19 unique; 16S primer: N=1 929, 1 178 unique). The species overlap between protocols for the 12S was only 1.2% (n=1), and for the 16S primer it was 3.5% (n=402).

**Figure 3:**
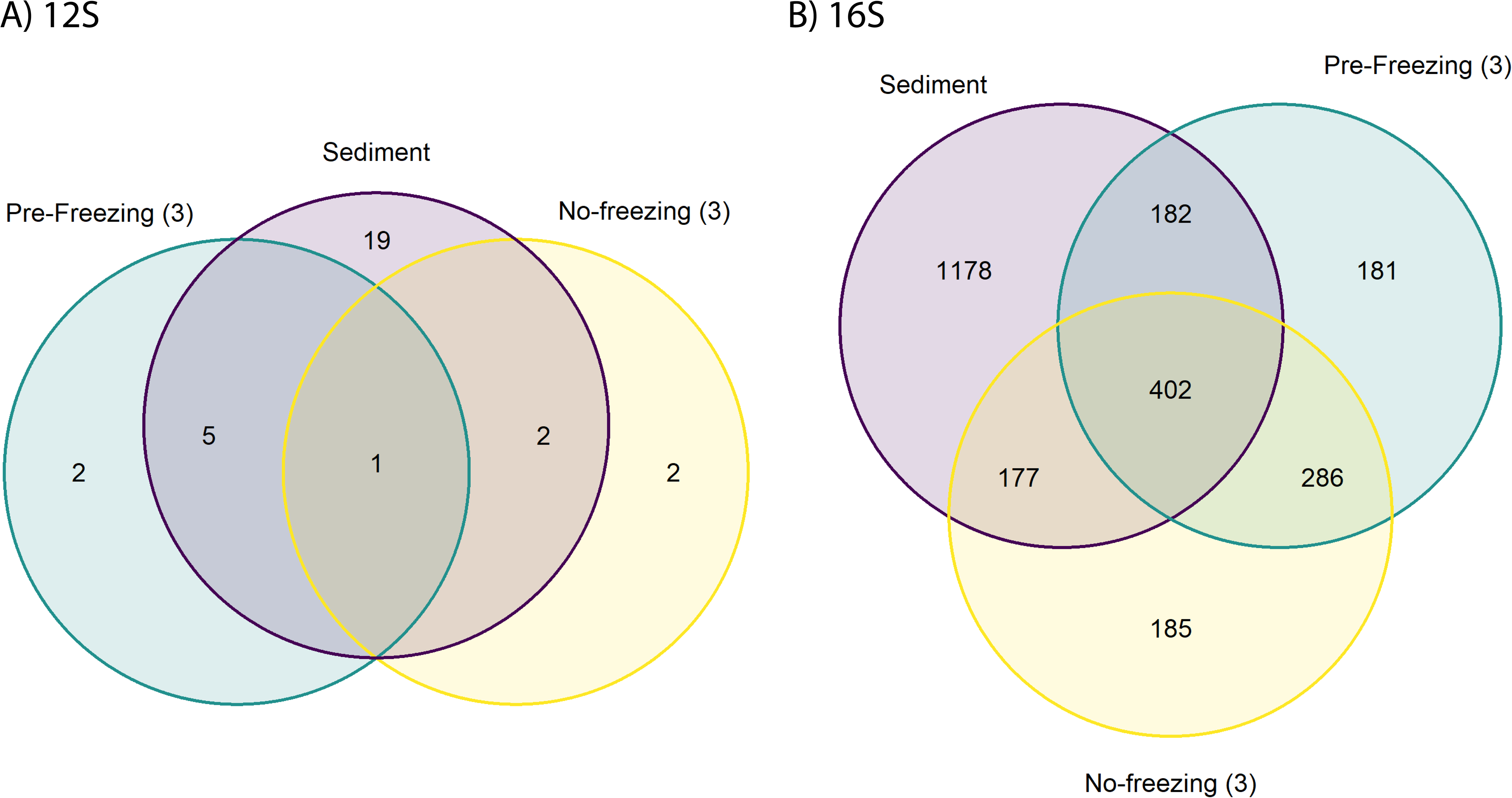
Venn diagrams of A) 12S and B) 16S primers showing the number of species found at and between each protocol. Sediment samples showed the highest number of unique species for both primers, although for the 12S dataset, about 28% are the result of contamination from tag-jumping.

Species rarefaction curves also show that the sediment protocol is further from reaching saturation compared to both filtration protocols, for both the 12S and 16S primers (Fig. 4), although there was more variation between the replicates for the 12S sediment protocol. For the 12S primer, there is a significant difference in the slope of the species curves between the sediment and no freezing protocols (Fig. 5, Table 2), while for the 16S, all pairwise comparisons showed significant differences.

**Figure 4:**
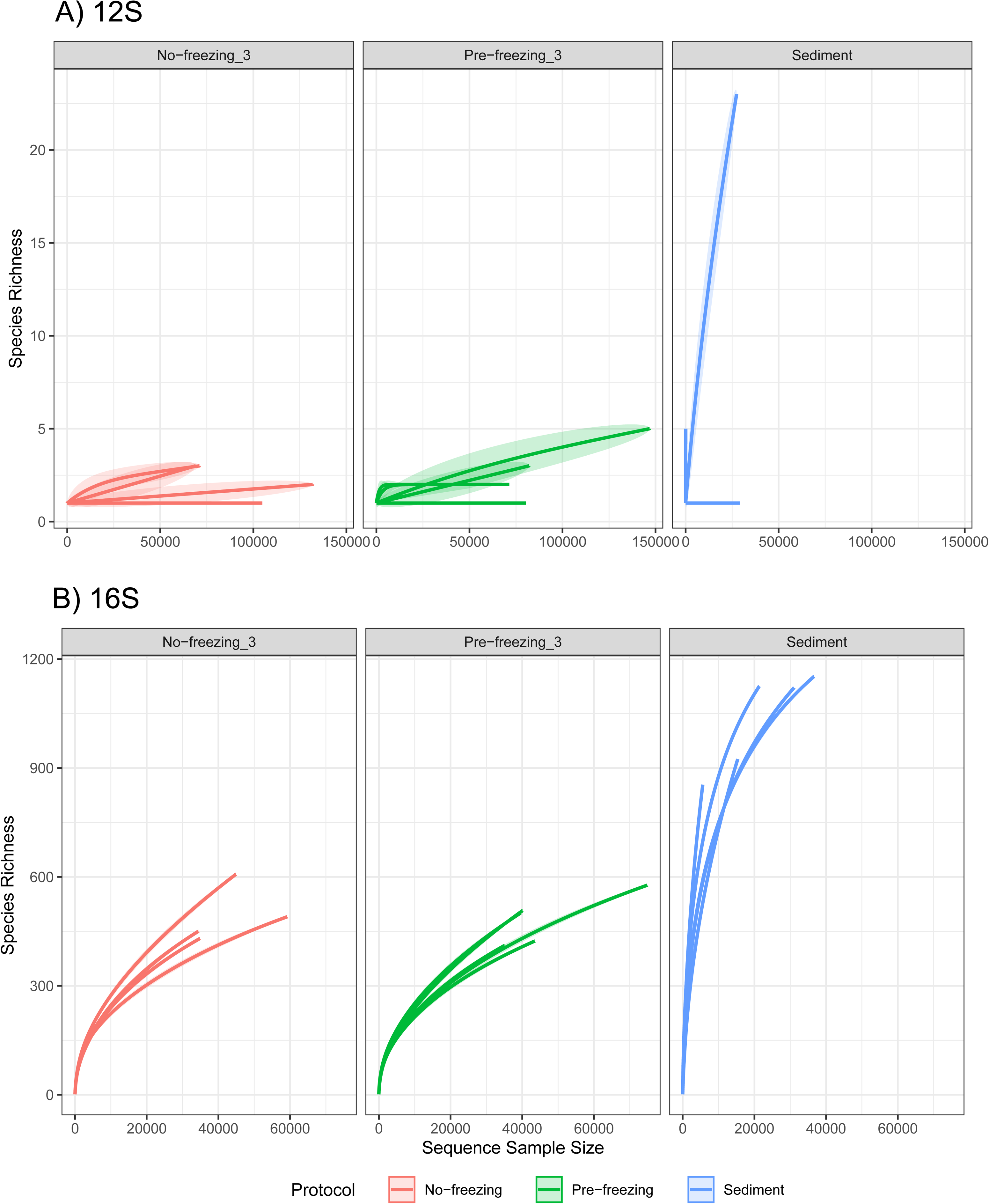
Species rarefaction curves based on sequencing effort for each protocol. A) 12S primer; B) 16S primer. With the exception of the water samples for the 12S primer, none of the curves have reached a plateau, although we expect the high diversity seen for the 12S sediment samples be due to contamination from tag-jumping.

**Figure 5:**
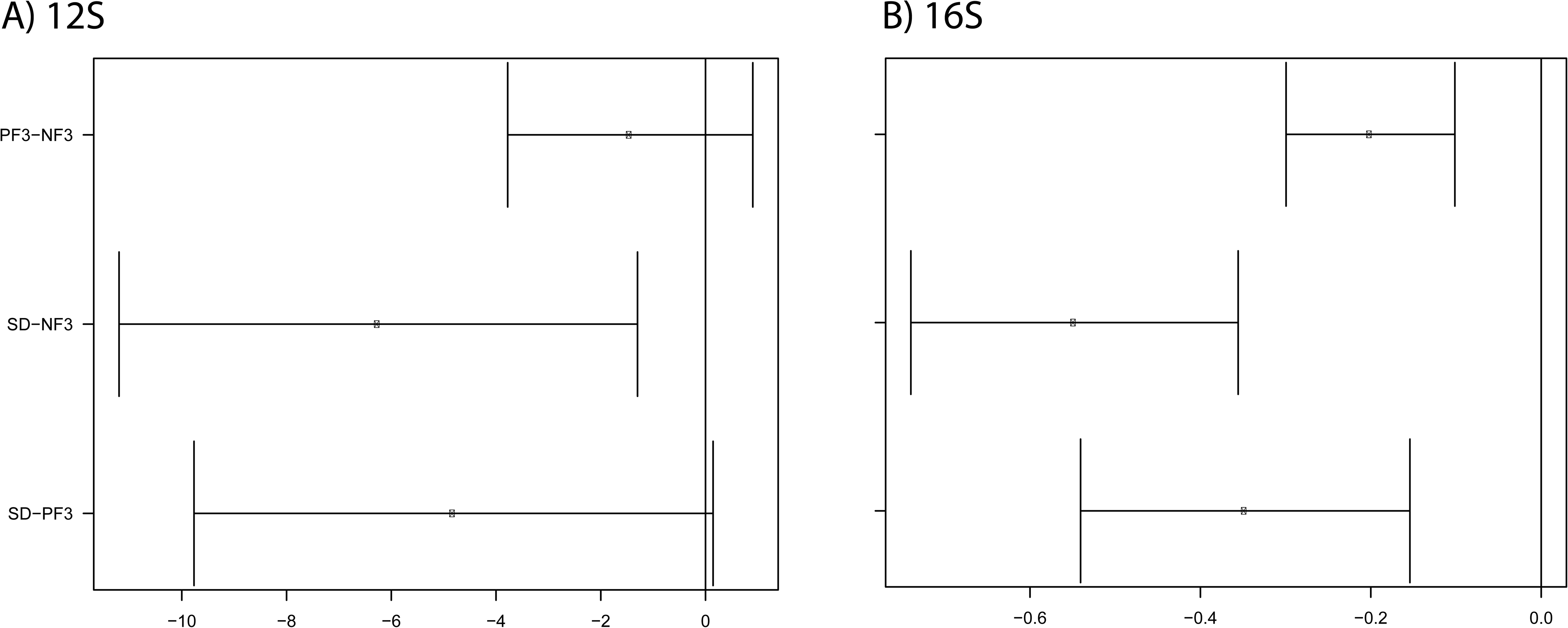
Confidence interval (CI) for slopes of rarefaction curves (Fig. 3) for each pairwise comparison of the different protocols. Only the comparison between pre-versus no freezing water samples, and pre-freezing versus sediment samples for the 12S primer (A) have come out non significant. The remaining comparisons showed significant differences between rarefaction slopes.

**Table 2:**
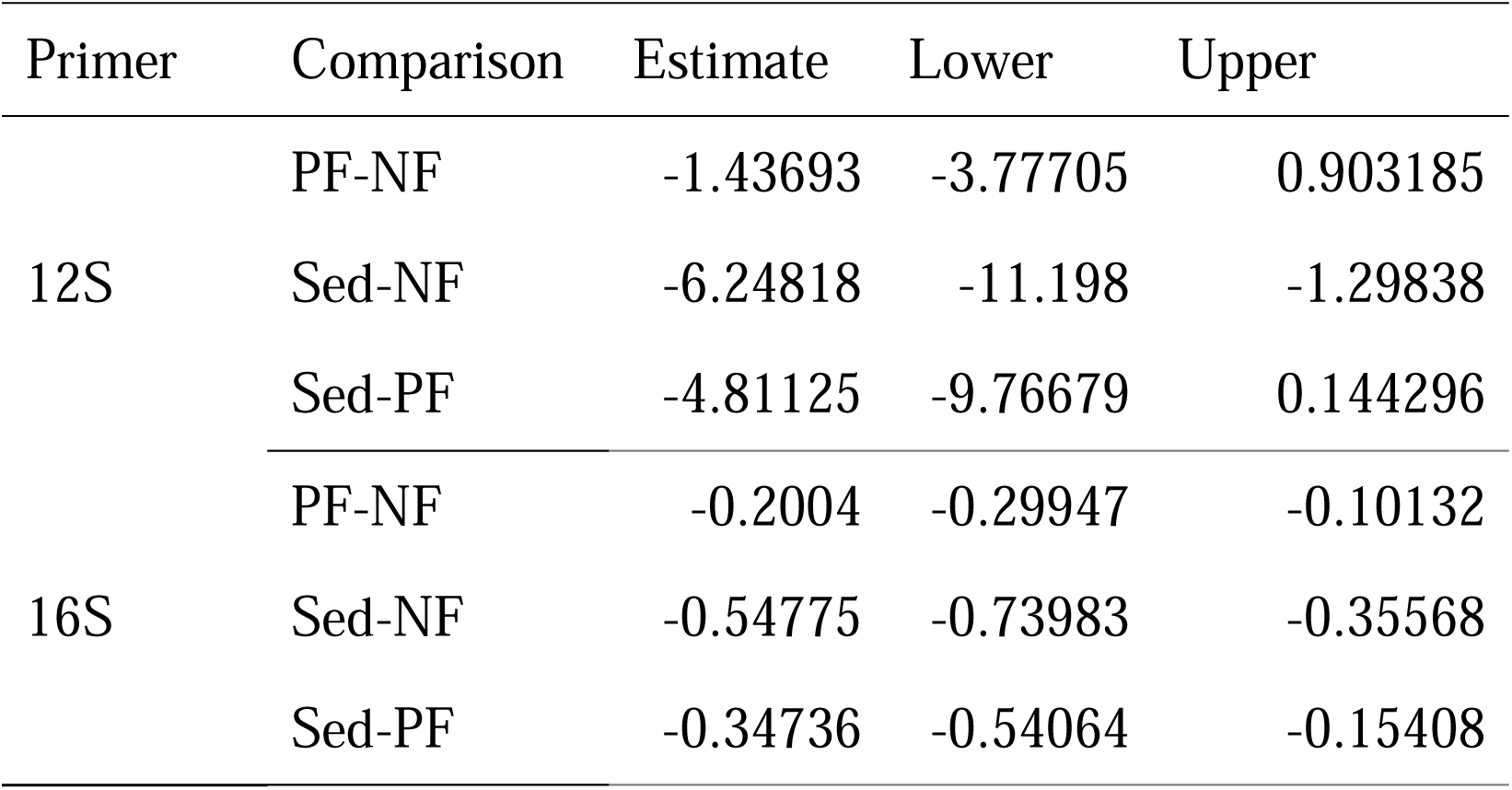
Confidence interval estimates and upper/lower limits of species rarefaction curves (Fig. 4). NF: no freezing; PF: pre-freezing; Sed: sediment.

### Differential Abundance

In the DESeq2 analysis, there was no significant difference between species abundance for the 12S primer in any of the protocols’ pairwise comparisons. In ALDEx2, the tidewater goby (*E. newberryi*) was the only species with an effect size > 1 in the comparisons of filtration protocols (NF and PF) against the sediment protocol, being underrepresented in the latter (Fig. 6). The proportion of overlap of the 95% CI of the effect size was not zero, but they were small nonetheless (NF: 0.010; PF: 0.014) (Table S4).

**Figure 6:**
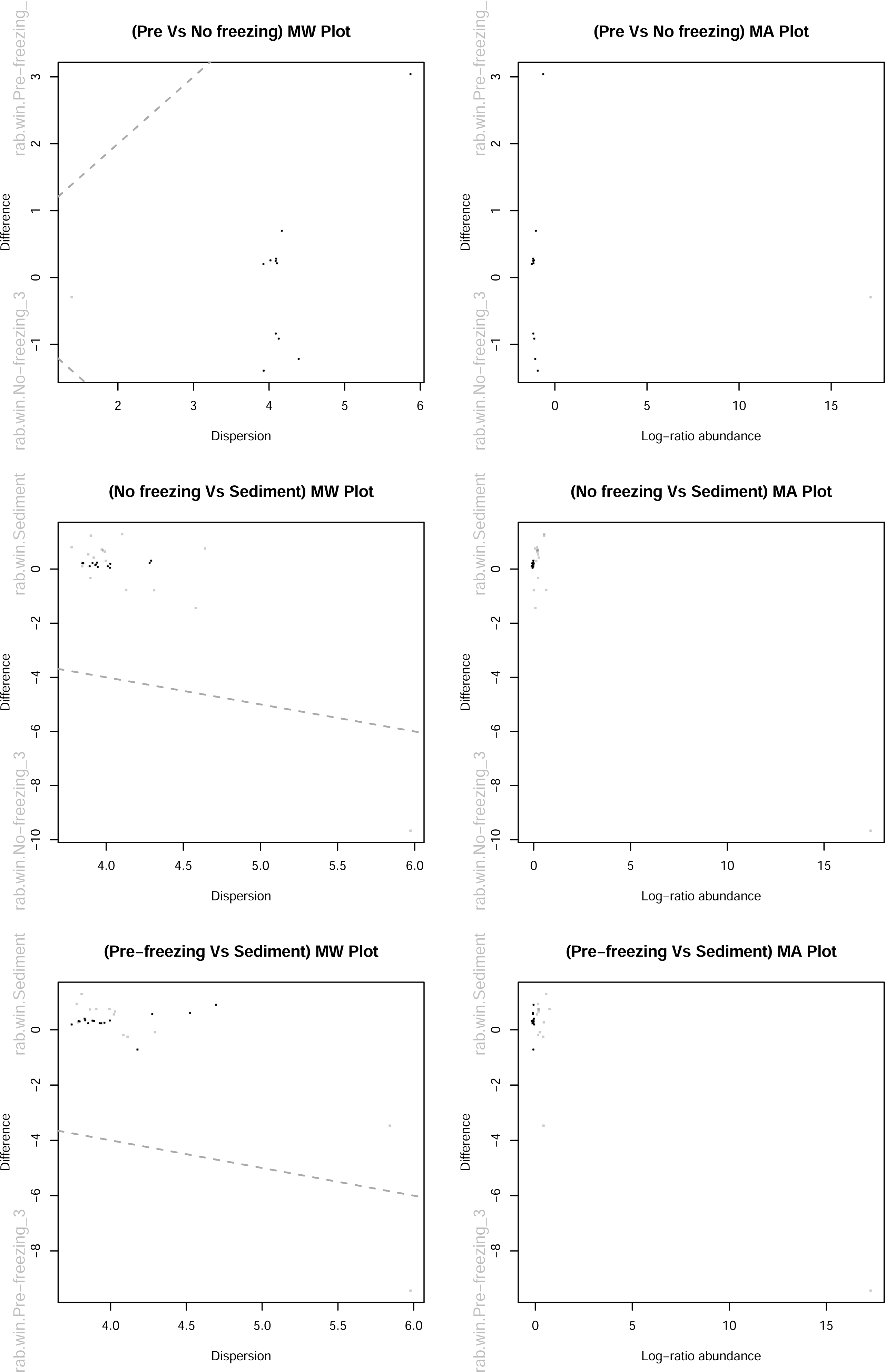
MW (left) and BA (right) plots for 12S primer dataset. In both plots, gray represents features that are abundant, but not differentially abundant; black are rare, but not differentially abundant. Diagonal dashed lines are shown for zero-intercept with slope of ± 1. Single grey point outside of intercept is the tidewater goby species (E. newberryi).

In the case of the 16S primer, the DESeq2 analysis showed no significant difference in comparison between the filtration protocols (NF versus PF). However, there were significant differences in the pairwise comparisons between filtration protocols and sediment protocol (NF vs. Sed; PF vs. Sed; Fig. 6, Tables S2–3). The five most differentially abundant species with highest abundance in the filtration protocols, relative to the sediment, were representatives of the families Aphanizomenonaceae, Comamonadaceae and Flavobacteriaceae (in both NF and PF comparisons); and Hemiselmidaceae and Geminigeraceae (PF protocol only). These comprise groups of cyanobacteria (Aphanizomenonaceae) and algae (Hemiselmidaceae and Geminigeraceae), as well as environmental bacteria (Comamonadaceae and Flavobacteriaceae).

The most differentially abundant species with highest abundance in the sediments were representatives of the families Catenulaceae, Fragilariaceae and an archaea assigned to the Thaumarchaeota phylum (in both NF and PF comparisons); plus Woeseiaceae and Elphidiidae (NF protocol only); and Anaerolineaceae and Desulfobacteraceae (PF protocol only). These comprise groups of diatoms (Catenulaceae and Fragilariaceae), environmental bacteria (Woeseiaceae, Anaerolineaceae and Desulfobacteraceae) and archaea (Thaumarchaeota), and foraminiferans (Elphidiidae).

In the ALDEx2 analysis, there were many species that showed an effect size > 1 (Fig. 8), but only a few showed proportions of overlap < 0.001 (none in the NF versus PF comparison) (Table S5). For the species that fall within these criteria (effect > 1, overlap < 0.001), in the comparisons between filtration protocols and sediment protocol (NF vs. Sed; PF vs. Sed), there were 61 species overrepresented in the NF protocol compared to 39 in the sediment; and 73 species overrepresented in the PF protocol compared to the sediment (there was no species overrepresented in the sediment that met the criteria above) (Table S5). Taxa overrepresented in both filtration protocols (NF and PF) relative to sediment are in the families Comamonadaceae, Flavobacteriaceae and Rhodobacteraceae. Both Comamonadaceae and Flavobacteriaceae were also represented in the DESeq2 results and comprise groups of environmental bacteria. Rhodobacteraceae are heterotrophic bacteria usually found in association with algae in marine environments (Bischoff et al., 2021). In the NF protocol, Cryomorphaceae is the most represented family, with seven species, and comprises aquatic bacteria generally found in locations rich in organic carbon (Bowman, 2014).

**Figure 7:**
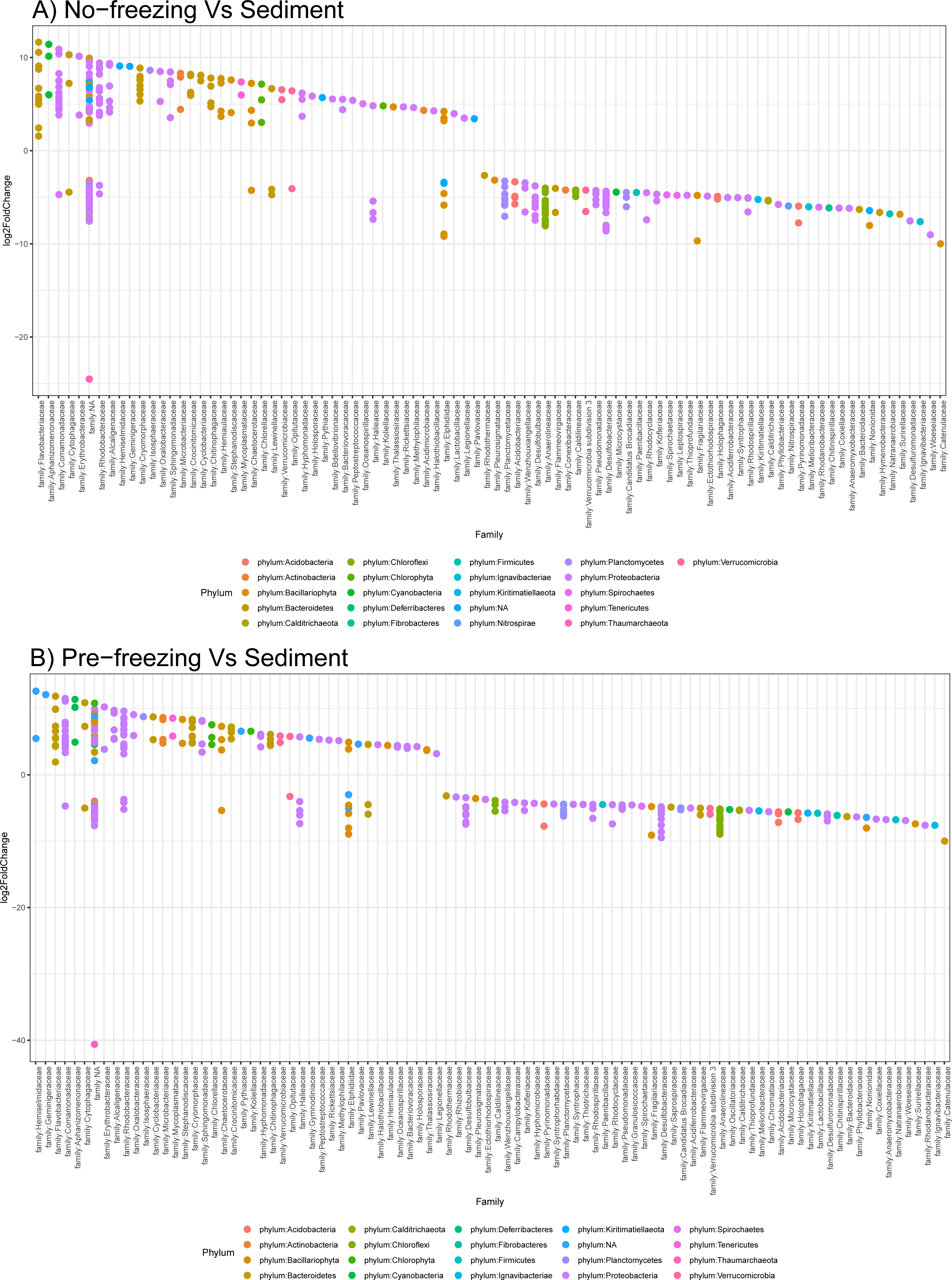
Plots of log2fold change of families of bacteria and archaea (16S primer) for the pairwise comparison between A) no freezing versus sediment; and B) pre-freezing versus sediment. Circles are colored by phylum. Species present above zero are overrepresented in the pre-or no freezing protocol, and species below the zero threshold are overrepresented in the sediments.

**Figure 8:**
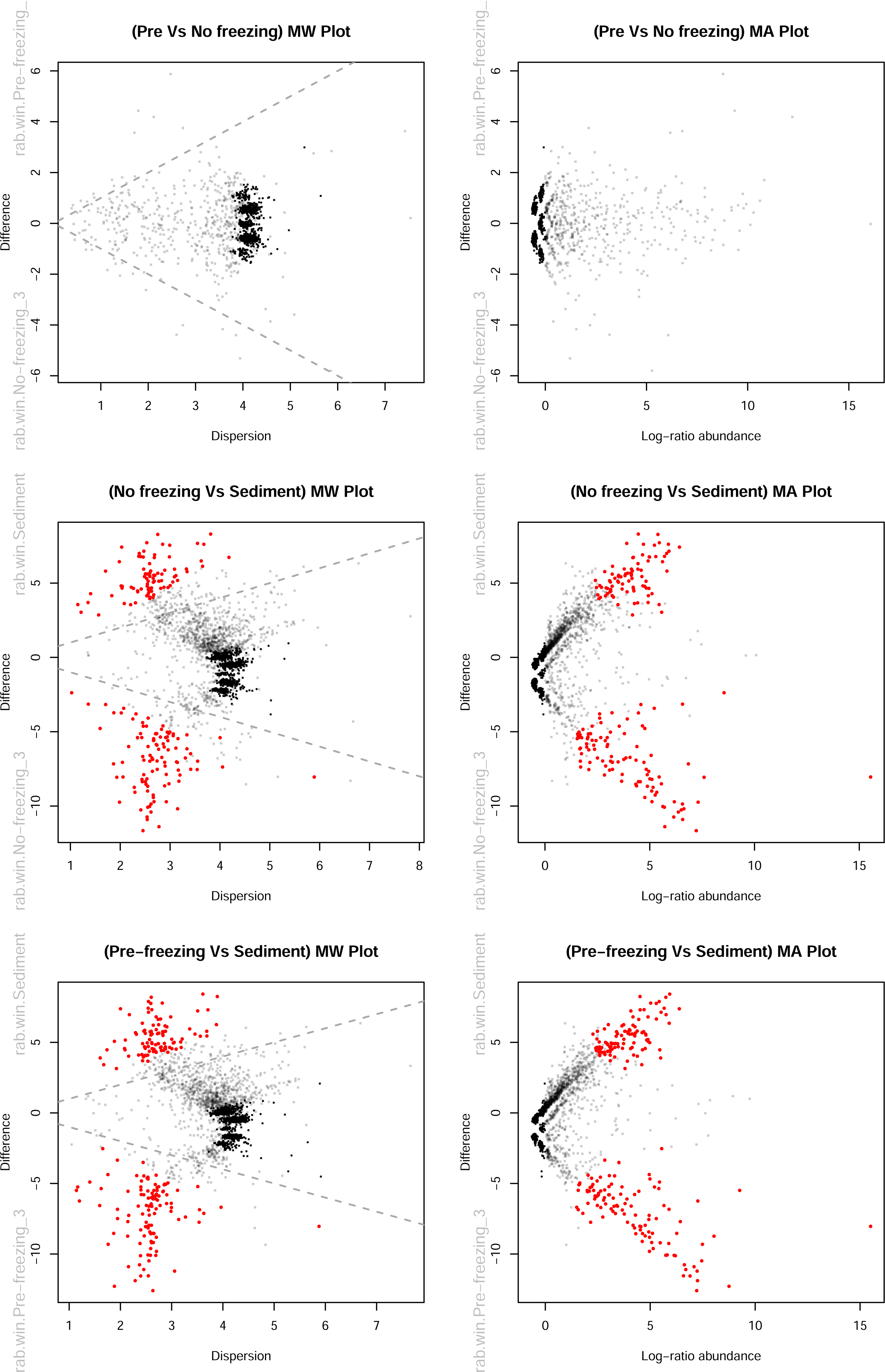
MW (left) and BA (right) plots for 16S primer dataset. In both plots, gray represents features that are abundant, but not differentially abundant; black are rare, but not differentially abundant. Red represents features called as differentially abundant with q < 0.1. Diagonal dashed lines are shown for zero-intercept with slope of ± 1.

In the sediments, when compared to the NF protocol, the most overrepresented family is the Anaerolineaceae with 12 species, followed by Desulfobacteraceae (5), Nonionidae (2) and Elphidiidae (2). Anaerolineaceae, Desulfobacteraceae and Elphidiidae were also found in the DESeq2 results. The first two comprise environmental bacteria, while the latter is a family of foraminiferans (such as the Noniodidae family).

### Beta diversity

For the rarefied dataset, there were no discernible differences in assemblage composition in the CAP analysis for the 12S primer (Fig. 9A). One sediment replicate is driving most of the difference (CAP1=86%) with the overrepresentation of many tropical species, likely tag-jump contaminants. The PERMANOVA result showed a relatively weak effect overall (R^2^ = 0.38; p = 0.05), with the pairwise test showing a relatively weak effect for the NF versus PF comparison (R^2^ = 0.20), followed by stronger effects in the sediment comparisons (NF vs. Sed: R^2^ = 0.41; PF vs. Sed: R^2^ = 0.31) (Table 3). The lack of significant differences between the filtration and sediment protocols could have been driven by the loss of three sediment replicates when rarefying the dataset.

**Table 3:** PERMANOVA and pairwise PERMANOVA (rarefied dataset) between all three protocols: no-freezing, pre-freezing and sediment. P.adjusted is the adjusted p-value after FDR correction.

| Primer | PERMANOVA |  | Pairwise PERMANOVA |  |  |  |  |
| --- | --- | --- | --- | --- | --- | --- | --- |
|  | R2 | Pr(>F) | Comparison | F.Model | R2 | p.value | p.adjusted |
| 12S | 0.3803 | 0.05 | No-freezing vs Pre-freezing | 2.07252 | 0.20576 | 0.169 | 0.295 |
|  |  |  | No-freezing vs Sediment | 3.56051 | 0.41592 | 0.295 | 0.295 |
|  |  |  | Pre-freezing vs Sediment | 2.25713 | 0.31102 | 0.229 | 0.295 |
| 16S | 0.6922 | 0.001 | No-freezing vs Pre-freezing | 10.3356 | 0.56369 | 0.006 | 0.009 |
|  |  |  | No-freezing vs Sediment | 12.1022 | 0.63355 | 0.007 | 0.009 |
|  |  |  | Pre-freezing vs Sediment | 12.5474 | 0.6419 | 0.009 | 0.009 |

When using the eDNA index, the CAP analysis for the 12S primer showed that many of the species driving the differences in assemblage composition were the tropical species that are coming from the tag-jumping contamination (Fig. S5A). For example, we see overrepresentation in the sediment protocol of Stegastes nigricans and Caranx melampygus; and in the NF protocol, Sphyraena barracuda. Nevertheless, we also see some other species that are known to be found in the lagoon, such as the Eucyclogobius newberryi, being mostly overrepresented in the filtration protocols (NF and PF) compared to the sediments; and Gila orcutii, overrepresented in the NF protocol. Two species of dubious taxonomic assignment are also overrepresented in the sediment: Phoxinus phoxinus (as discussed in the previous ‘Diversity’ section); and Acanthogobius flavimanus, which is a species of goby native to Asia, but that has been recorded previously in California estuaries (Nico et al., 2022). The PERMANOVA result showed a weaker effect than in the rarefied dataset (R2 = 0.21; p = 0.055), including for the pairwise comparisons (Table 4).

**Table 4:** Pairwise PERMANOVA and pairwise PERMANOVA (eDNA index dataset) between all three protocols: no-freezing, pre-freezing and sediment pre-and no freezing water prior to filtration and sediment samples. P.adjusted is the adjusted p-value after FDR correction.

| Primer | PERMANOVA |  | Pairwise PERMANOVA |  |  |  |  |
| --- | --- | --- | --- | --- | --- | --- | --- |
|  | R2 | Pr(>F) | Comparison | F.Model | R2 | p.value | p.adjusted |
| 12S | 0.2113 | 0.055 | No-freezing vs Pre-freezing | 1.25387 | 0.1355 | 0.297 | 0.297 |
|  |  |  | No-freezing vs Sediment | 1.76131 | 0.18044 | 0.112 | 0.168 |
|  |  |  | Pre-freezing vs Sediment | 1.69357 | 0.17471 | 0.077 | 0.168 |
| 16S | 0.4417 | 0.001 | No-freezing vs Pre-freezing | 1.47905 | 0.15603 | 0.005 | 0.0135 |
|  |  |  | No-freezing vs Sediment | 5.96537 | 0.42715 | 0.017 | 0.017 |
|  |  |  | Pre-freezing vs Sediment | 6.51459 | 0.44883 | 0.009 | 0.0135 |

For the rarefied 16S primer dataset, the different protocols showed discernible differences in assemblage composition in the CAP analysis. The first axis explains most of the total variation (CAP1=96%), with the tidewater goby being the most underrepresented in the sediment compared to the filtration protocols (NF and PF), especially in the NF protocol (Fig. 9B). The sediment protocol was also slightly overrepresented by a few other species. One of them was identified as *Nitrosopelagicus brevis*, a species of uncultured “Candidatus”, ammonia-oxidizing archaea (Thaumarchaeota) found mainly in the epi- and upper mesopelagic environments of the open oceans (Santoro et al., 2015). There are also two species of *Monomorphina*, (*M. pyrum* and *M. pseudonordstedti*) that belong to the Euglenaceae family, a group of eukaryotic flagellates found in freshwater environments. Lastly, there is *Elphidium williamsoni*, a foraminifera belonging to the family Elphidiidae found in tidal flats of the North Sea. CAP2 identifies the variation (14%) differentiating the filtration protocols, with the most distinguishing species being the *Guillardia theta*, a species of flagellate algae belonging to the family Geminigeraceae, overrepresented in the PF protocol. Most of these species were also recovered in the DESeq2 and ALDEx2 analyses, with the exception of both Monomorphina species. The PERMANOVA result showed a relatively large effect (R^2^ = 0.69; p = 0.001), as well as for all the pairwise comparisons (R^2^ > 0.5; Table 3).

The species represented in the rarefied dataset differ from the ones found when using the eDNA index for the 16S primer. Most of the community assemblage difference in the eDNA index (CAP1=85%) is driven by differences between filtration and sediment protocols, with six species being underrepresented in the latter: *Burkholderiales bacterium* TP637, *Curvibacter* sp. UKPF8, *Diaphorobacter ruginosibacter* and *Verminephrobacter aporrectodeae* (Comamonadaceae); *beta proteobacterium Mzo1* (Oxalobacteraceae); and *Stella humosa* (*Peptostreptococcaceae*). Most of these species were also recovered in the DESeq2 and ALDEx2 analyses, with the exception of *V. aporrectodeae*. The PERMANOVA result also showed relatively high effect (R^2^ = 0.44; p = 0.001), but for the pairwise comparisons the effect between the filtration protocols (NF vs. PF) was relatively small (R2 = 0.15;Table 4).

**Figure 9:**
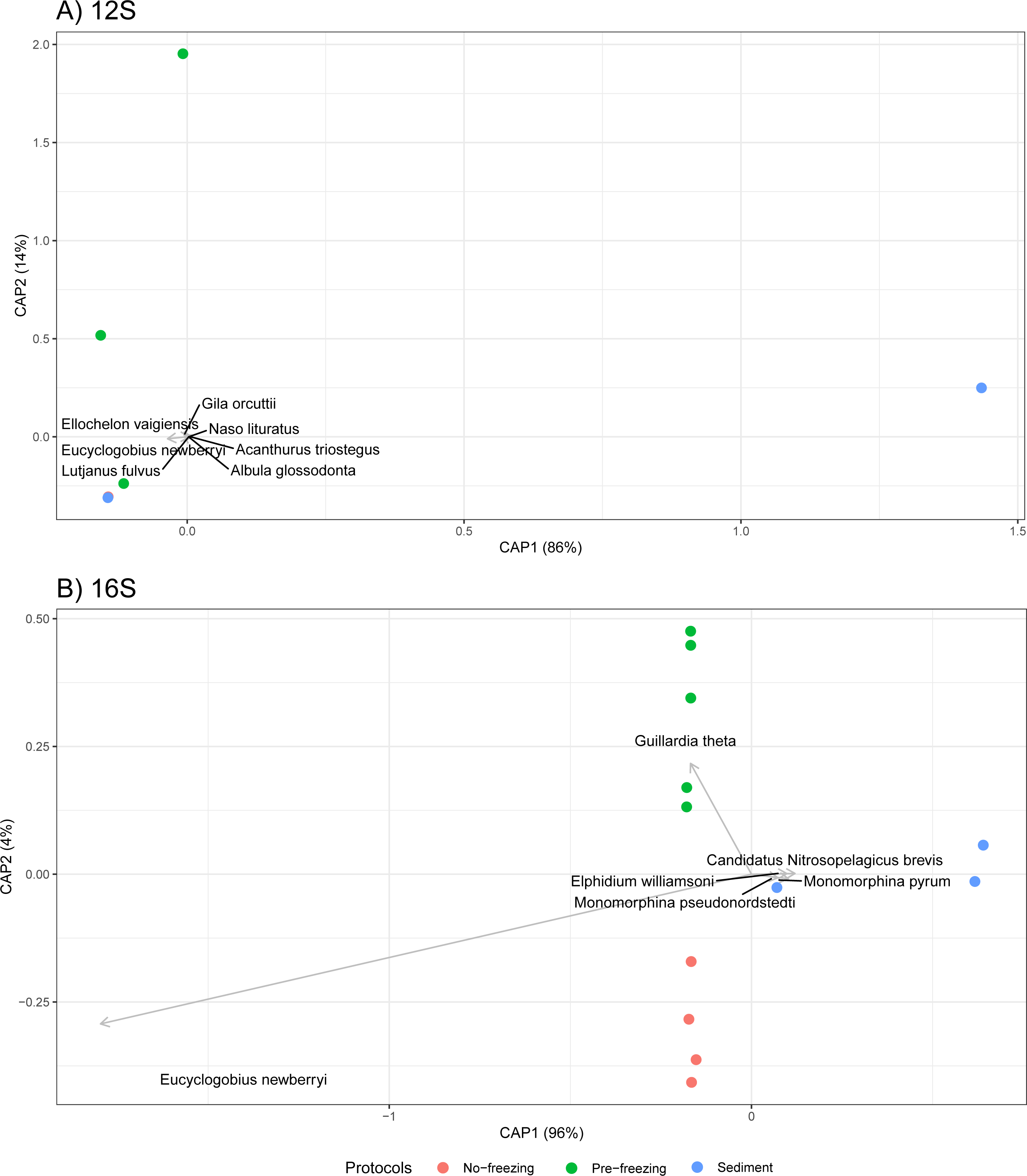
Constrained Analysis of Principal Coordinates (CAP) of rarefied A) 12S and B) 16S primer datasets. Circles are colored by protocol. While there were no discernible differences in assemblage composition for the 12S primer, in the 16S primer *Guillardia theta* is overrepresented in the PF protocol in comparison to the NF protocol. *Eucyclogobius newberyii* is underrepresented in the sediments compared to the filtration protocols, especially the NF protocol.

### Discussion

Standardized protocols to process eDNA are under development (e.g. Bohmann et al., 2021), but to implement these efficiently it is necessary to compare biases in taxa detection associated with different protocols. Here, we have explored the detection biases in community composition introduced by freezing water samples prior to filtration (for storage purposes), and the use of sediment samples as an alternative to sampling turbid waters. We find that pre-freezing water does incur some effect on the recovery of community composition, but most strongly for the 16S primer, while having a relatively small effect for the 12S when using larger, cellulose nitrate pore size filters (3 μm).The sediment protocol was able to recover eDNA from organisms that inhabit the water column, however, due to high variability in read abundance among replicates, we suggest increasing the number of biological replicates and/or the volume sampled in the field.

### Tag-jumping contamination

Contamination concerns are usually centered around pre-sequencing, during the field and wet laboratory work. These are of fundamental importance and care should be taken by sterilizing equipment and using negative controls. However, previous literature shows that the sequencing phase can be another source of contamination, generating up to 10% of contaminated reads by tag-jumping (Larsson et al., 2018; Schnell et al., 2015), which can skew analyses of taxa abundance and composition towards the rare taxa. There are ways to help minimize this issue by making use of dual indexing (Kircher et al., 2012)—although see Caroe and Bohmann (2020) for a library approach without dual indexing—, and amplification of positive controls. The latter can be used to track the rate and level of contamination after sequencing to guide read cutoffs on samples (Deiner et al., 2017; Port et al., 2016).

### Bioinformatics and data pre-processing

We relied on a bioinformatic approach developed by the metabaR package, adapted from Esling et al. (2015), to reduce the issue of contamination from tag-jumping, since it does not rely solely on the use of positive controls (which we lacked in this analysis) to make the estimated cutoff thresholds. However, after manually checking the fish dataset (12S primer), the final taxonomic tables still contained reads assigned to taxa that are not found in coastal lagoons in California (Table S1). Some of it might be contamination from tag-jumping, although we cannot rule out the possibility that for a few of these species the eDNA could have come from local aquaria, as some are known in the pet trade (e.g. *Acanthurus achilles*). We also cannot disregard the limitations of the reference database, especially related to the absence of estuarine and lagoonal taxa that may lead to dubious assignments to non-local related species (Nagarajan et al., 2022). Due to inability to completely remove potential tag-jump contaminants from the dataset, we can expect a bias towards the rare taxa that will inflate diversity metrics in our samples for all primer sets.

The sediment protocol generally showed higher variability among replicates compared to filtration protocols for both primer sets, both in number of reads and community composition (Fig. S3-4). The greater consistency of filtration replicates is a result of the single source for the water samples (the large jug). This means that potentially, multiple biological replicates could be sampled at the site and pooled for fewer downstream analysis, optimizing cost and labor (Sato et al., 2017; Dickie et al., 2018), however, since this was not the focus of this work this design would require further investigation. Sediment replicates were done by individually sampling the bottom of the lagoon. Although replicates were done a few centimeters apart, the bottom of the lagoon appears to have small-scale heterogeneity. The spatial variation of soil and sediment samples is recognized in the literature (Perkins et al., 2014; Taberlet, Prud’Homme, et al., 2012), and can be caused by sediment composition but also by the flow dynamic and distribution of eDNA in the water column. While this variability has been shown to occur for water samples as well in lentic environments (Harper et al., 2019 and references therein), the heterogeneity of water replicates in this system still requires further investigation.

The sediment protocol also had an overall lower number of reads compared to the filtration protocols for both primer sets (Fig. S3). The lower number of reads seems to go against the expectations that eDNA can be more concentrated in sediments (Dell’Anno & Corinaldesi, 2004; Harper et al., 2019; Turner et al., 2015). This could be due to a few issues, some of which may interact. First, it could be related to a faster degradation and/or turn-over rates of eDNA in the sediment, which are determined by the soil and eDNA characteristics, as well as enzymatic and microbial activities (Levy-Booth et al., 2007; Pietramellara et al., 2009; Torti et al., 2015). The overall lower abundance of eDNA in the sediments could also be driven by increased inhibition (Buxton et al., 2017; Pawlowski et al., 2022). Even though we used a specific soil extraction kit for both sediment and filtration protocols, the purification steps in the protocol could still not have been enough to reduce inhibition in the sediment as well as for the water samples. Lastly, this could have been driven by the much lower volume of sediment used: 0.25-0.3 g versus 500 mL for water samples.

There is also the fact that this type of environment is affected by scouring (purging of sediment to the ocean) during high precipitation events and increased flow of freshwater. However, since the sediment collection was done out of the rainy season and the lagoon was closed by a sandbar with no signs of scouring, we are confident that this was not a factor that could have caused the decreased ability to recover eDNA from the sediments. Therefore, we expect that this difference in read abundance between sediment and water samples would be more related to the other factors mentioned above, such as eDNA degradation and turn-over rates, inhibition, and different processed volumes. Considering both the high variability and the lower sequencing throughput of the sediment replicates, we advise using a modified sampling protocol, e.g. the one developed by Taberlet, Prud’Homme, et al. (2012) that includes increasing the number of replicates and mixing larger volumes before downstream processing.

### Diversity

Considering that contamination through tag-jumping could be inflating the numbers of rare species in the dataset, the steepness and lack of a plateau for many of the species rarefaction curves could be artificial (Fig. 4). This is especially evident for the 12S primer since we were able to manually investigate the taxonomy tables (Table S1). However, this lack of a plateau is an expected outcome from environmental samples (Alberdi et al., 2018), and has been shown to occur more acutely in a coastal lagoon in California when compared to other environments in California (Shirazi et al., 2021)—albeit the authors were looking specifically at plants and fungi. The high number of species recovered from the sediment for the 16S primer (Fig. 3) is likely driven by the recovery of a rich and complex sediment biota that is not paralleled in the water column.

The low taxonomic assignment to the species level for some of the dubious fish species found in our dataset, e.g *Phoxinus phoxinus, Odontesthes* spp. And *Sebastes pachycephalus*, also highlight the need to expand barcoding efforts to the local estuarine taxa to improve reference databases. On the other hand, *Fundulus diaphanus*, the northeastern killifish, did receive a few high taxonomic scores at the species level, which merit further consideration for biomonitoring of coastal lagoons in the region.

Pre-freezing water prior to filtration had an effect on the species curves of the 16S primer dataset, but not so much on the 12S. This could be explained by how differently eDNA molecules are found in the environment for these two different groups of organisms, and how freezing and thawing water would impact them. In the case of the fish fauna, the DNA that is shed from the organisms would be either found within cells, or adsorbed to colloids (Liang & Keeley, 2013; Torti et al., 2015; Turner et al., 2014). Even if cell walls were to disintegrate from the freezing and thawing process, they could still release intact mitochondria (which range from 1-8 μm in length) that could still be captured by our 3 μm pore size filters. On the other hand, bacteria and archaea, which are prokaryotic and often single celled organisms, would have their DNA released directly to the medium and pass through the larger pore size filters (>0.2 μm). Nevertheless, this freezing effect on cell walls has been shown to not always occur and likely be species-dependent (Sekar et al., 2009; Suomalainen et al., 2006).

### Differential abundance

Pre-freezing water did introduce bias in species abundance compared to the no freezing protocol, although the effect was distinctively larger for the 16S than for the 12S primer dataset when using larger, cellulose nitrate pore size filters (3 μm). Our results seem to align to other reports, where it was shown that freezing had differential effects on detection and relative abundance of different prokaryotic taxa (Kwambana et al., 2011; Sekar et al., 2009; Suomalainen et al., 2006). The smaller effect seen for the 12S dataset could be related to the different qualities and properties of eDNA in the environment, as mentioned in the previous section (‘Diversity’). Larger, mitochondrial molecules from Eukaryotes may be more easily captured in the filters compared to the smaller celled, prokaryotic organisms targeted by the 16S primer.

Naturally, due to eDNA precipitation and resuspension, we expect to capture some community overlap between water and surficial sediment samples, however abundances should be different as consequence of the origin and fate of the eDNA in the environment and the processes acting on it throughout (Torti et al., 2015). As expected, in the differential abundance analyses, we see overrepresentation of families of algae (Hemiselmidaceae and Geminigeraceae), environmental and aquatic bacteria (Comamonadaceae, Flavobacteriaceae, Cryomorphaceae and Rhodobacteraceae), and cyanobacteria (Aphanizomenonaceae) in the water samples (NF and PF protocols).There is also significantly higher representation of presumptively benthic diatoms (Catenulaceae and Fragilariaceae), environmental bacteria (Anaerolineaceae and Desulfobacteraceae) and foraminiferans (Elphidiidae and Nonionidae) in the sediment. In addition, the types of environmental bacteria most abundant in the sediments were typical of soil and sediments elsewhere. Of particular note are those from anoxic environments (e.g. Anaerolineaceae and Desulfobacteraceae) as lagoon sediments in this region are often dark and sulfide-rich, consistent with anoxia and sufler-cycling.

In the family Flavobacteriaceae there are important pathogens of fish and humans that belong to the genus *Flavobacterium*. Suomalainen et al. (2006) found that *F. columnare* was more susceptible to having its cell walls disrupted to freezing due to high amounts of DNAases, lyases and proteases, likely connected to its pathogenicity, which then led to lower rates of DNA recovery. The species found in our dataset was *F. johnsoniae*, a species not known to be pathogenic–albeit with low species taxonomic score. Given that there was no difference in abundance for this species in our NF and PF protocols, which is in contrast with the results for the pathogenic species, *F. columnare*, this might relate to a true non-pathogenic organism. However, considering that the endangered northern tidewater goby often achieves high abundance in this lagoon, more detailed assessment of the *Flavobacterium* species inhabiting this site would be of interest.

The other species assignment that draws our attention is the archea Candidatus *Nitrosopelagicus brevis* (Thaumarchaeota), which was shown as more abundant in sediment than water samples. As mentioned earlier, this is a pelagic species, normally found in the open ocean worldwide. Although coastal lagoons are subject to marine input, the relatively high concentration in sediment is unexpected and merits inquiry, especially considering that the confidence in its taxonomic assignment was low across reads. More likely, this represents a new environmental archaea that is abundant in coastal lagoon sediments.

### Beta diversity

McMurdie and Holmes (2014) recommends against rarefying datasets due to the risk of removing true, rare ASVs. However, in our case, where we were unable to completely remove tag-jumping contaminants, this pre-process could help alleviate some of the noise caused by contaminants. Nevertheless, the CAP and PERMANOVA results on both the rarefied and standardized (eDNA index) dataset are mostly in agreement, and show greater differences in assemblage composition for the 16S primer than for the 12S primer, especially between the filtration and sediment protocols (NF vs. Sed; PF vs. Sed). Although results from the 12S primer were highly affected by the tag-jump contaminants in the dataset standardized using the eDNA index (Fig. S5A), the CAP analysis was able to capture the overrepresentation of tidewater goby in the filtration protocols compared to the sediment, a signal that was lost in the rarefied data (Fig. 9A).

For the rarefied 16S primer dataset, most species that were over- and underrepresented in the CAP analysis were also captured by the differential abundance analyses, such as *Guillardia theta* (Geminigeraceae), which was overrepresented in the PF protocol compared to the NF protocol. In addition, the species of foraminifera, *Elphidium williamsoni* (Elphidiidae) and the archea Candidatus *Nitrosopelagicus brevis* (Thaumarchaeota) were found to be overrepresented in the sediment compared to the filtration protocols. The CAP results on the dataset standardized using the eDNA index (Fig. S5B) showed different species as underrepresented in the sediment compared to the filtration protocols, but those also showed up as significantly different in the differential abundance analyses, with the exception of one, *Verminephrobacter aporrectodeae*.

Interestingly, the CAP analysis was also able to capture the underrepresentation of tidewater gobies (*E. newberryi*) in the sediment protocol on the 16S primer when compared to the NF protocol (Fig. 9B). The 16S ribosomal RNA region have been used for fish species detection, and is not uncommon that different primer sets are taxonomically congruent (Shu, Lidwig & Peng, 2020; Alberdi et al., 2018). This reinforces the idea discussed earlier (‘Bioinformatics and data pre-processing’ section) that fish eDNA, at least in this environment, is less concentrated in the sediment than in the water column, which contradicts other findings from the literature (Perkins et al., 2014; Turner et al., 2015). But it is worth noting that this underrepresentation of fish eDNA in the sediment was found to be of small effect for the 12S primer, though, and there could be some bias related to how these two genes behave and degrade differently in the environment for the fish fauna.

### Lessons Learned

Here is a list of recommendations and best practices for eDNA sampling and analysis in coastal environments that we have learned throughout this work and believe will be useful for others working in similar environments with turbid water and highly heterogeneous sediment/soil:

1. Filtered water samples had an overall higher number of reads compared to sediment for both primer sets. Therefore, we recommend the use of this protocol as it will increase chances of species detection;
2. If using sediment samples, we recommend increasing the number of replicates and mixing larger volumes before processing for DNA extractions (as in Taberlet, Prud’Homme, et al., 2012);
3. Pre-freezing water samples prior to filtration can be an effective storage solution, at least for Eukaryotes, and when used in combination with cellulose nitrate filters of up to 3 μm pore size filters;
4. The use of dual-indexing and positive controls during library preparation will help minimize and address cross-contamination from tag-jumping, as is now widely recognized in many best-practice protocols (e.g. Deiner et al., 2017; Goldberg et al., 2016);
5. Although rarefying the dataset is not recommended (McMurdie & Holmes, 2014), we recognize that it can aid in reducing the noise of contaminants from your dataset, as long as they are rare. Otherwise, the use of eDNA index (Kelly et al., 2019) can be an alternative to standardize your dataset.

## Conclusions

In this work, we assessed environmental DNA protocols for use in coastal lagoons, a highly dynamic habitat at the intersection of terrestrial, freshwater and marine environments. Pre-freezing water combined with the use of larger pore size filters (at least up to 3 μm) can be a viable alternative for storage and processing of turbid water samples, at least in the case of fish communities (12S, MiFish). However, the use of sediment samples as an alternative to processing water samples should be done with caution, and at minimum the number of biological replicates and/or volume sampled should be increased. Also, while sediment samples were able to recover eDNA from organisms commonly found in the water column, such as the tidewater goby, this was achieved during a period of relatively long lagoon closure, when there was no recent scouring of sediments to the ocean.

While we expect these guidelines to be helpful in the development of strategies to use eDNA as a monitoring resource in similar environments, protocol testing is still strongly advised whenever possible, especially when working in a new system. Much work is necessary to understand the full potential eDNA brings for the conservation and restoration of endangered species and habitats.

## Acknowledgements

Funding was provided by the National Council for Scientific and Technological Development of Brazil (Rachel Turba) under Grant No. 209261/2014-5 and by NOAA Sea Grant 120651698:1. Funding for the CALeDNA sample processing, infrastructure, and personnel was provided by the University of California Research Initiatives (UCRI) Catalyst grant CA-16-376437 and Howard Hughes Medical Institute (HHMI) Professors Grant GT10483. We are very grateful for all the help provided by the CALeDNA team, but would like to give special thanks to Teia Schweizer, who personally trained us in the bench work. Huge thanks to Ryan Kelly, for helping streamline the design and analysis of this paper and for always being so responsive via email, and to reviewer David Murray-Stoker for a much detailed and comprehensive feedback that helped improve this manuscript.

## References

Alberdi, A., Aizpurua, O., Gilbert, M. T. P., & Bohmann, K. (2018). Scrutinizing key steps for reliable metabarcoding of environmental samples. Methods in Ecology and Evolution, 9(1), 134–147. https://doi.org/10.1111/2041-210X.12849

Ballard, J., Pezda, J., Spencer, D., & Plantinga, A. (n.d.). *An Economic Valuation of Southern California Coastal Wetlands*. http://scwrp.org/wp-content/uploads/2017/06/SoCalWetlands_FinalReport.pdf

Bischoff, V., Zucker, F., & Moraru, C. (2021). Marine Bacteriophages. In D. H. Bamford & M. Zuckerman (Eds.), Encyclopedia of Virology (Fourth Edition) (pp. 322–341). Academic Press. https://doi.org/10.1016/B978-0-12-809633-8.20988-6

Bohmann, K., Chua, P., Holman, L. E., & Lynggaard, C. (2021). DNAqua-Net conference unites participants from around the world with the quest to standardize and implement DNA-based aquatic biomonitoring. Environmental DNA, 3(5), 884–888. https://doi.org/10.1002/edn3.207

Bowman, J. P. (2014). The Family Cryomorphaceae. In E. Rosenberg, E. F. DeLong, S. Lory, E. Stackebrandt, & F. Thompson (Eds.), *The Prokaryotes* (pp. 539–550). Springer Berlin Heidelberg. https://doi.org/10.1007/978-3-642-38954-2_135

Buxton, A. S., Groombridge, J. J., & Griffiths, R. A. (2017). Is the detection of aquatic environmental DNA influenced by substrate type? PLOS ONE, 12(8), e0183371. https://doi.org/10.1371/journal.pone.0183371

Callahan, B. J., McMurdie, P. J., Rosen, M. J., Han, A. W., Johnson, A. J. A., & Holmes, S. P. (2016). DADA2: High-resolution sample inference from Illumina amplicon data. Nature Methods, 13(7), 581–583. https://doi.org/10.1038/nmeth.3869

Caporaso, J. G., Lauber, C. L., Walters, W. A., Berg-Lyons, D., Huntley, J., Fierer, N., Owens, S. M., Betley, J., Fraser, L., Bauer, M., Gormley, N., Gilbert, J. A., Smith, G., & Knight, R. (2012). Ultra-high-throughput microbial community analysis on the Illumina HiSeq and MiSeq platforms. The ISME Journal, 6(8), 1621–1624. https://doi.org/10.1038/ismej.2012.8

Caroe, C., & Bohmann, K. (2020). Tagsteady: A metabarcoding library preparation protocol to avoid false assignment of sequences to samples. *BioRxiv*.

Chen, H. (2018). VennDiagram: Generate High-Resolution Venn and Euler Plots (1.6.20). https://CRAN.R-project.org/package=VennDiagram

Curd, E., Gomer, J., Kandlikar, G., Gold, Z., Ogden, M., & Shi, B. (2018). *The Anacapa Toolkit*. https://github.com/limey-bean/Anacapa

Curd, E., Kandlikar, G., & Gomer, J. (2018). *CRUX: Creating Reference libraries Using eXisting tools*. https://github.com/limey-bean/CRUX_Creating-Reference-libraries-Using-eXisting-tools

Deiner, K., Bik, H. M., Mächler, E., Seymour, M., Lacoursière-Roussel, A., Altermatt, F., Creer, S., Bista, I., Lodge, D. M., & De Vere, N. (2017). Environmental DNA metabarcoding: Transforming how we survey animal and plant communities. Molecular Ecology, 26(21), 5872–5895.

Deiner, K., Walser, J.-C., Mächler, E., & Altermatt, F. (2015). Choice of capture and extraction methods affect detection of freshwater biodiversity from environmental DNA. Biological Conservation, 183, 53–63.

Dejean, T., Valentini, A., Miquel, C., Taberlet, P., Bellemain, E., & Miaud, C. (2012). Improved detection of an alien invasive species through environmental DNA barcoding: The example of the American bullfrog *Lithobates catesbeianus*: *Alien invasive species detection using eDNA*. Journal of Applied Ecology, 49(4), 953–959. https://doi.org/10.1111/j.1365-2664.2012.02171.x

Dell’Anno, A., & Corinaldesi, C. (2004). Degradation and Turnover of Extracellular DNA in Marine Sediments: Ecological and Methodological Considerations Degradation and Turnover of Extracellular DNA in Marine Sediments: Ecological and Methodological Considerations. Applied and Environmental Microbiology, 70(7), 4384–4386. https://doi.org/10.1128/AEM.70.7.4384

Doi, H., Uchii, K., Matsuhashi, S., Takahara, T., Yamanaka, H., & Minamoto, T. (2017). Isopropanol precipitation method for collecting fish environmental DNA. Limnology and Oceanography: Methods, 15(2), 212–218. https://doi.org/10.1002/lom3.10161

Earl, D. A., Louie, K. D., Bardeleben, C., Swift, C. C., & Jacobs, D. K. (2010). Rangewide microsatellite phylogeography of the endangered tidewater goby, a genetically subdivided coastal fish with limited marine dispersal. Conservation Genetics, 11, 103–104. https://doi.org/10.1007/s10592-009-0008-9

Esling, P., Lejzerowicz, F., & Pawlowski, J. (2015). Accurate multiplexing and filtering for high-throughput amplicon-sequencing. Nucleic Acids Research, 43(5), 2513–2524. https://doi.org/10.1093/nar/gkv107

*FASTX-Toolkit*. (n.d.). Retrieved January 11, 2018, from http://hannonlab.cshl.edu/fastx_toolkit/

Fernandes, A. D., Macklaim, J. M., Linn, T. G., Reid, G., & Gloor, G. B. (2013). ANOVA-Like Differential Expression (ALDEx) Analysis for Mixed Population RNA-Seq. PLoS ONE, 8(7), e67019. https://doi.org/10.1371/journal.pone.0067019

Ficetola, G. F., Miaud, C., Pompanon, F., & Taberlet, P. (2008). Species detection using environmental DNA from water samples. Biology Letters, 4(4), 423–425. https://doi.org/10.1098/rsbl.2008.0118

Gao, X., Lin, H., Revanna, K., & Dong, Q. (2017). A Bayesian taxonomic classification method for 16S rRNA gene sequences with improved species-level accuracy. BMC Bioinformatics, 18(1), 247. https://doi.org/10.1186/s12859-017-1670-4

Gbif.Org. (2022). Occurrence Download (p. 170487) [Darwin Core Archive]. The Global Biodiversity Information Facility. https://doi.org/10.15468/DL.HTJ3HT

Goldberg, C. S., Turner, C. R., Deiner, K., Klymus, K. E., Thomsen, P. F., Murphy, M. A., Spear, S. F., McKee, A., Oyler-McCance, S. J., & Cornman, R. S. (2016). Critical considerations for the application of environmental DNA methods to detect aquatic species. Methods in Ecology and Evolution, 7(11), 1299–1307.

Harper, L. R., Buxton, A. S., Rees, H. C., Bruce, K., Brys, R., Halfmaerten, D., Read, D. S., Watson, H. V., Sayer, C. D., & Jones, E. P. (2019). Prospects and challenges of environmental DNA (eDNA) monitoring in freshwater ponds. Hydrobiologia, 826(1), 25–41.

Hinlo, R., Gleeson, D., Lintermans, M., & Furlan, E. (2017). Methods to maximise recovery of environmental DNA from water samples. PloS One, 12(6), e0179251.

Jacobs, D. K., Stein, E. D., & Longcore, T. (2011). Classification of California Estuaries Based on Natural Closure Patterns: Templates for Restoration and Management Management. Technical Report, *August*, 1–72.

Kandlikar, G. (2020). ranacapa: Utility Functions and “shiny” App for Simple Environmental DNA Visualizations and Analyses (0.1.0). https://github.com/gauravsk/ranacapa

Kelly, R. P., Shelton, A. O., & Gallego, R. (2019). Understanding PCR Processes to Draw Meaningful Conclusions from Environmental DNA Studies. Scientific Reports, 9(1), 12133. https://doi.org/10.1038/s41598-019-48546-x

Kircher, M., Sawyer, S., & Meyer, M. (2012). Double indexing overcomes inaccuracies in multiplex sequencing on the Illumina platform. Nucleic Acids Research, 40(1), e3–e3. https://doi.org/10.1093/nar/gkr771

Kumar, G., Farrell, E., Reaume, A. M., Eble, J. A., & Gaither, M. R. (2022). One size does not fit all: Tuning eDNA protocols for high- and low-turbidity water sampling. Environmental DNA, 4(1), 167–180. https://doi.org/10.1002/edn3.235

Kwambana, B. A., Mohammed, N. I., Jeffries, D., Barer, M., Adegbola, R. A., & Antonio, M. (2011). Differential effects of frozen storage on the molecular detection of bacterial taxa that inhabit the nasopharynx. BMC Clinical Pathology, 11(1), 2. https://doi.org/10.1186/1472-6890-11-2

Langmead, B., & Salzberg, S. L. (2012). Fast gapped-read alignment with Bowtie 2. Nature Methods, 9(4), 357–359. https://doi.org/10.1038/nmeth.1923

Laramie, M. B., Pilliod, D. S., Goldberg, C. S., & Strickler, K. M. (2015). Environmental DNA sampling protocol—Filtering water to capture DNA from aquatic organisms. *U.S Geological Survey Techniques and Methods*, Book 2(Chapter A13), 15 p. https://doi.org/10.3133/TM2A13

Larsson, A. J. M., Stanley, G., Sinha, R., Weissman, I. L., & Sandberg, R. (2018). Computational correction of index switching in multiplexed sequencing libraries. Nature Methods, 15(5), 305–307. https://doi.org/10.1038/nmeth.4666

Leray, M., Yang, J. Y., Meyer, C. P., Mills, S. C., Agudelo, N., Ranwez, V., Boehm, J. T., & Machida, R. J. (2013). A new versatile primer set targeting a short fragment of the mitochondrial COI region for metabarcoding metazoan diversity: Application for characterizing coral reef fish gut contents. Frontiers in Zoology, 10(1), 34. https://doi.org/10.1186/1742-9994-10-34

Levy-Booth, D. J., Campbell, R. G., Gulden, R. H., Hart, M. M., Powell, J. R., Klironomos, J. N., Pauls, K. P., Swanton, C. J., Trevors, J. T., & Dunfield, K. E. (2007). Cycling of extracellular DNA in the soil environment. Soil Biology and Biochemistry, 39(12), 2977– 2991. https://doi.org/10.1016/j.soilbio.2007.06.020

Li, J., Lawson Handley, L.-J., Read, D. S., & Hänfling, B. (2018). The effect of filtration method on the efficiency of environmental DNA capture and quantification via metabarcoding. Molecular Ecology Resources, 18(5), 1102–1114.

Liang, Z., & Keeley, A. (2013). Filtration Recovery of Extracellular DNA from Environmental Water Samples. Environmental Science & Technology, 47(16), 9324–9331. https://doi.org/10.1021/es401342b

Love, M. I., Huber, W., & Anders, S. (2014). Moderated estimation of fold change and dispersion for RNA-seq data with DESeq2. Genome Biology, 15(12), 550. https://doi.org/10.1186/s13059-014-0550-8

Majaneva, M., Diserud, O. H., Eagle, S. H., Boström, E., Hajibabaei, M., & Ekrem, T. (2018). Environmental DNA filtration techniques affect recovered biodiversity. Scientific Reports, 8(1), 1–11.

Martin, M. (2013). Cutadapt removes adapter sequences from high-throughput sequencing reads. EMBnet.Journal, 17(1), 10–12. http://journal.embnet.org/index.php/embnetjournal/article/view/200/479

McMurdie, P. J., & Holmes, S. (2013). phyloseq: An R Package for Reproducible Interactive Analysis and Graphics of Microbiome Census Data. PLoS ONE, 8(4), e61217. https://doi.org/10.1371/journal.pone.0061217

McMurdie, P. J., & Holmes, S. (2014). Waste Not, Want Not: Why Rarefying Microbiome Data Is Inadmissible. PLoS Computational Biology, 10(4), e1003531. https://doi.org/10.1371/journal.pcbi.1003531

Miya, M., Sato, Y., Fukunaga, T., Sado, T., Poulsen, J. Y., Sato, K., Minamoto, T., Yamamoto, S., Yamanaka, H., Araki, H., Kondoh, M., & Iwasaki, W. (2015). MiFish, a set of universal PCR primers for metabarcoding environmental DNA from fishes: Detection of more than 230 subtropical marine species. Royal Society Open Science, 2(7). http://rsos.royalsocietypublishing.org/content/2/7/150088.abstract

Nagarajan, R. P., Bedwell, M., Holmes, A. E., Sanches, T., Acuña, S., Baerwald, M., Barnes, M. A., Blankenship, S., Connon, R. E., Deiner, K., Gille, D., Goldberg, C. S., Hunter, M. E., Jerde, C. L., Luikart, G., Meyer, R. S., Watts, A., & Schreier, A. (2022). Environmental DNA Methods for Ecological Monitoring and Biodiversity Assessment in Estuaries. Estuaries and Coasts, 45(7), 2254–2273. https://doi.org/10.1007/s12237-022-01080-y

Oksanen, J., Blanchet, F. G., Friendly, M., Kindt, R., Legendre, P., McGlinn, D., Minchin, P. R., O’Hara, R. B., Simpson, G. L., Solymos, P., Stevens, M. H. H., Szoecs, E., & Wagner, H. (2019). vegan: Community Ecology Package (2.5-6). https://CRAN.R-project.org/package=vegan

Pawlowski, J., Bruce, K., Panksep, K., Aguirre, F. I., Amalfitano, S., Apothéloz-Perret-Gentil, L., Baussant, T., Bouchez, A., Carugati, L., Cermakova, K., Cordier, T., Corinaldesi, C., Costa, F. O., Danovaro, R., Dell’Anno, A., Duarte, S., Eisendle, U., Ferrari, B. J. D., Frontalini, F., … Fazi, S. (2022). Environmental DNA metabarcoding for benthic monitoring: A review of sediment sampling and DNA extraction methods. Science of The Total Environment, 818, 151783. https://doi.org/10.1016/j.scitotenv.2021.151783

Perkins, T. L., Clements, K., Baas, J. H., Jago, C. F., Jones, D. L., Malham, S. K., & McDonald, J. E. (2014). Sediment Composition Influences Spatial Variation in the Abundance of Human Pathogen Indicator Bacteria within an Estuarine Environment. PLoS ONE, 9(11), e112951. https://doi.org/10.1371/journal.pone.0112951

Pietramellara, G., Ascher, J., Borgogni, F., Ceccherini, M. T., Guerri, G., & Nannipieri, P. (2009). Extracellular DNA in soil and sediment: Fate and ecological relevance. Biology and Fertility of Soils, 45(3), 219–235. https://doi.org/10.1007/s00374-008-0345-8

Pilliod, D. S., Goldberg, C. S., Arkle, R. S., Waits, L. P., & Richardson, J. (2013). Estimating occupancy and abundance of stream amphibians using environmental DNA from filtered water samples. Canadian Journal of Fisheries and Aquatic Sciences, 70(8), 1123–1130. https://doi.org/10.1139/cjfas-2013-0047

Port, J. A., O’Donnell, J. L., Romero-Maraccini, O. C., Leary, P. R., Litvin, S. Y., Nickols, K. J., Yamahara, K. M., & Kelly, R. P. (2016). Assessing vertebrate biodiversity in a kelp forest ecosystem using environmental DNA. Molecular Ecology, 25(2), 527–541. https://doi.org/10.1111/mec.13481

R Core Team. (2018). R: A language and environment for statistical computing. R Foundation for Statistical Computing. https://www.R-project.org/

Rees, H. C., Maddison, B. C., Middleditch, D. J., Patmore, J. R. M., & Gough, K. C. (2014). REVIEW: The detection of aquatic animal species using environmental DNA – a review of eDNA as a survey tool in ecology. Journal of Applied Ecology, 51(5), 1450–1459. https://doi.org/10.1111/1365-2664.12306

Robson, H. L. A., Noble, T. H., Saunders, R. J., Robson, S. K. A., Burrows, D. W., & Jerry, D. R. (2016). Fine-tuning for the tropics: Application of eDNA technology for invasive fish detection in tropical freshwater ecosystems. Molecular Ecology Resources, 16(4), 922– 932. https://doi.org/10.1111/1755-0998.12505

RStudio Team. (2020). RStudio: Integrated Development for R. RStudio, PBC. http://www.rstudio.com/

Sales, N. G., Wangensteen, O. S., Carvalho, D. C., & Mariani, S. (2019). Influence of preservation methods, sample medium and sampling time on eDNA recovery in a neotropical river. Environmental DNA, 1(2), edn3.14. https://doi.org/10.1002/edn3.14

Santoro, A. E., Dupont, C. L., Richter, R. A., Craig, M. T., Carini, P., McIlvin, M. R., Yang, Y., Orsi, W. D., Moran, D. M., & Saito, M. A. (2015). Genomic and proteomic characterization of “Candidatus Nitrosopelagicus brevis”: An ammonia-oxidizing archaeon from the open ocean. Proceedings of the National Academy of Sciences, 112(4), 1173–1178. https://doi.org/10.1073/pnas.1416223112

Sard, N. M., Herbst, S. J., Nathan, L., Uhrig, G., Kanefsky, J., Robinson, J. D., & Scribner, K. T. (2019). Comparison of fish detections, community diversity, and relative abundance using environmental DNA metabarcoding and traditional gears. Environmental DNA, 1(4), 368–384. https://doi.org/10.1002/edn3.38

Schaarschmidt, F., & Gerhard, D. (2019). PairwiseCI: Confidence Intervals for Two Sample Comparisons (0.1-27). https://CRAN.R-project.org/package=pairwiseCI

Schnell, I. B., Bohmann, K., & Gilbert, M. T. P. (2015). Tag jumps illuminated—Reducing sequence-to-sample misidentifications in metabarcoding studies. Molecular Ecology Resources, 15(6), 1289–1303. https://doi.org/10.1111/1755-0998.12402

SCWRP. (2018). Wetlands on the Edge: The Future of Southern California’s Wetlands: Regional Strategy 2018 (p. 142). California State Coastal Conservancy. scwrp.databasin.org

Sekar, R., Kaczmarsky, L. T., & Richardson, L. L. (2009). Effect of Freezing on PCR Amplification of 16S rRNA Genes from Microbes Associated with Black Band Disease of Corals. Applied and Environmental Microbiology, 75(8), 2581–2584. https://doi.org/10.1128/AEM.01500-08

Shaffer, H. B., Fellers, G. M., Randal Voss, S., Oliver, J. C., & Pauly, G. B. (2004). Species boundaries, phylogeography and conservation genetics of the red-legged frog (Rana aurora/draytonii) complex. Molecular Ecology, 13(9), 2667–2677. https://doi.org/10.1111/j.1365-294X.2004.02285.x

Shirazi, S., Meyer, R. S., & Shapiro, B. (2021). Revisiting the effect of PCR replication and sequencing depth on biodiversity metrics in environmental DNA metabarcoding. Ecology and Evolution, 11(22), 15766–15779. https://doi.org/10.1002/ece3.8239

Smart, A. S., Weeks, A. R., Rooyen, A. R., Moore, A., McCarthy, M. A., & Tingley, R. (2016). Assessing the cost-efficiency of environmental DNA sampling. Methods in Ecology and Evolution, 7(11), 1291–1298. https://doi.org/10.1111/2041-210X.12598

Stein, E. D., Cayce, K., Salomon, M., Bram, D. L., De Mello, D., Grossinger, R., & Dark, S. (2014). *Wetlands of the Southern California Coast: Historical Extent and Change Over Time* (SFEI Report 720; SCCWRP Technical Report 826; p. 58). Southern California Coastal Water Research Project and San Francisco Estuary Institute. https://www.caltsheets.org/socal/download.html

Suomalainen, L.-R., Reunanen, H., Ijäs, R., Valtonen, E. T., & Tiirola, M. (2006). Freezing Induces Biased Results in the Molecular Detection of *Flavobacterium columnare*. Applied and Environmental Microbiology, 72(2), 1702–1704. https://doi.org/10.1128/AEM.72.2.1702-1704.2006

Swift, C. C., Haglund, T. R., Ruiz, M., & Fisher, R. N. (1993). The Status and Distribution of the Freshwater Fishes of Southern California. Bulletin of the Southern California Academy of Sciences, 92(3), 101–167.

Swift, C. C., Spies, B., Ellingson, R. A., & Jacobs, D. K. (2016). A New Species of the Bay Goby Genus Eucyclogobius, Endemic to Southern California: Evolution, Conservation, and Decline. PloS One, 11(7), e0158543. https://doi.org/10.1371/journal.pone.0158543

Taberlet, P., Coissac, E., Pompanon, F., Brochmann, C., & Willerslev, E. (2012). Towards next-generation biodiversity assessment using DNA metabarcoding: NEXT-GENERATION DNA METABARCODING. Molecular Ecology, 21(8), 2045–2050. https://doi.org/10.1111/j.1365-294X.2012.05470.x

Taberlet, P., Prud’Homme, S. M., Campione, E., Roy, J., Miquel, C., Shehzad, W., Gielly, L., Rioux, D., Choler, P., Clément, J.-C., Melodelima, C., Pompanon, F., & Coissac, E. (2012). Soil sampling and isolation of extracellular DNA from large amount of starting material suitable for metabarcoding studies: EXTRACTION OF EXTRACELLULAR DNA FROM SOIL. Molecular Ecology, 21(8), 1816–1820. https://doi.org/10.1111/j.1365-294X.2011.05317.x

Takahara, T., Minamoto, T., & Doi, H. (2015). Effects of sample processing on the detection rate of environmental DNA from the Common Carp (Cyprinus carpio). Biological Conservation, 183, 64–69. https://doi.org/10.1016/j.biocon.2014.11.014

Thomsen, P. F., & Willerslev, E. (2015). Environmental DNA – An emerging tool in conservation for monitoring past and present biodiversity. Biological Conservation, 183, 4–18. https://doi.org/10.1016/j.biocon.2014.11.019

Torti, A., Lever, M. A., & Jørgensen, B. B. (2015). Origin, dynamics, and implications of extracellular DNA pools in marine sediments. Marine Genomics, 24, 185–196. https://doi.org/10.1016/j.margen.2015.08.007

Tsuji, S., Takahara, T., Doi, H., Shibata, N., & Yamanaka, H. (2019). The detection of aquatic macroorganisms using environmental DNA analysis—A review of methods for collection, extraction, and detection. Environmental DNA, 1(2), 99–108. https://doi.org/10.1002/edn3.21

Turba, R., Thai, G., & Jacobs, D. (2023). Different approaches to processing environmental DNA samples in turbid waters have distinct effects for fish, bacterial and archaea communities (Version 2, p. 2137292766 bytes) [Data set]. Dryad. https://doi.org/10.5068/D1BQ25

Turner, C. R., Barnes, M. A., Xu, C. C. Y., Jones, S. E., Jerde, C. L., & Lodge, D. M. (2014). Particle size distribution and optimal capture of aqueous macrobial eDNA. Methods in Ecology and Evolution, 5(7), 676–684. https://doi.org/10.1111/2041-210X.12206

Turner, C. R., Uy, K. L., & Everhart, R. C. (2015). Fish environmental DNA is more concentrated in aquatic sediments than surface water. Biological Conservation, 183, 93–102. https://doi.org/10.1016/j.biocon.2014.11.017

van der Loos, L. M., & Nijland, R. (2021). Biases in bulk: DNA metabarcoding of marine communities and the methodology involved. Molecular Ecology, 30(13), 3270–3288. https://doi.org/10.1111/mec.15592

WeatherSpark.com. (n.d.). Historical Weather Summer 2018 at Point Mugu Naval Air Warfare Center. WeatherSpark.Com. Retrieved June 8, 2022, from https://weatherspark.com/h/s/145310/2018/1/Historical-Weather-Summer-2018-at-Point-Mugu-Naval-Air-Warfare-Center;-California;-United-States#Figures-Rainfall

Williams, K. E., Huyvaert, K. P., & Piaggio, A. J. (2017). Clearing muddied waters: Capture of environmental DNA from turbid waters. PLOS ONE, 12(7), e0179282. https://doi.org/10.1371/journal.pone.0179282

Zinger, L., Lionnet, C., Benoiston, A.-S., Donald, J., Mercier, C., & Boyer, F. (2020). *metabaR: An R package for the evaluation and improvement of DNA metabarcoding data quality* [Preprint]. Bioinformatics. https://doi.org/10.1101/2020.08.28.271817

