## Supplemental Material for "Different approaches to processing environmental DNA samples in turbid waters have distinct effects for fish, bacterial and archaea communities"

Excluded dataset

For the sake of clarity and conciseness, we decided to exclude from the main paper the sequencing results from the centrifugation protocol and the second stage filtration using the 0.45 µm pore size filter. In a preliminary investigation, the substantially lower sequence throughput from these protocols reduced the power of the combined rarefied dataset. Besides, our assumption is that the bias driven by the water freezing protocol would still be captured by the first stage filtration. Some preliminary results of the centrifugation and second stage filtration protocols are shown in Figure S2, and the available data is provided at (<https://gitlab.com/rturba/coastal-lagoon-edna>).

CO1 Sequencing

It is hard to pinpoint exactly what was the issue that led to the substantive lower throughput of the CO1 primer compared to the other two. By following the CALeDNA protocol, PCR products were cleaned using magnetic beads, checked in agarose gel for size range, and insert size was taken into account when pooling libraries. Therefore, considering that we saw no bias at the quality check step when filtering reads, some error must have occurred when doing the bench work that affected all samples for this specific primer. We suspect it could have happened when calculating the cleaned PCR product concentrations using the fluorometer plate reader, since the CO1 primer was the only one that was read in a separate plate from the 12S and 16S primers.

Supplemental Figures and Tables

Table S1: List of fish species for the 12S primer dataset after the decontamination pipeline. Gbif = species listed in the GBIF database (Gbif.Org, 2022); lab_col = species listed in the laboratory collection database; tax_level = taxonomic level match between the eDNA and the other species database.

Table S2: List of species with significant differential abundance in the DESeq2 pairwise comparison between no freezing (NF) and sediment (Sed) protocols.

Table S3: List of species with significant differential abundance in the DESeq2 pairwise comparison between pre-freezing (PF) and sediment (Sed) protocols.

Table S4: 12S primer ALDEx2 taxonomy output from the t-test between: PF and NF protocols; NF and Sed protocols; and PF and Sed protocols.

Table S5: 16S primer ALDEx2 taxonomy output from the t-test between: PF and NF protocols; NF and Sed protocols; and PF and Sed protocols.

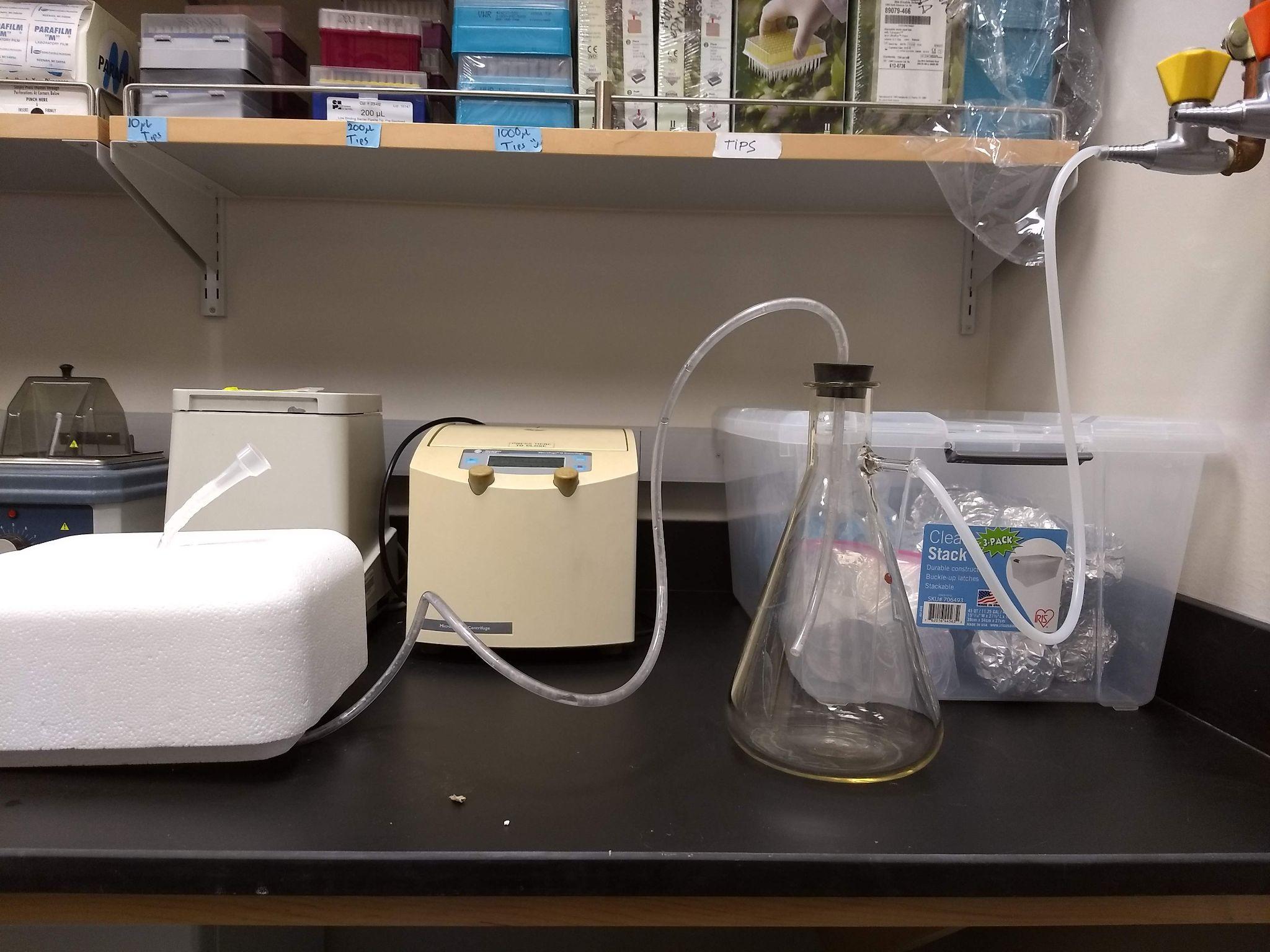

Figure S1: Adapted vacuum pump in the pre-PCR room.

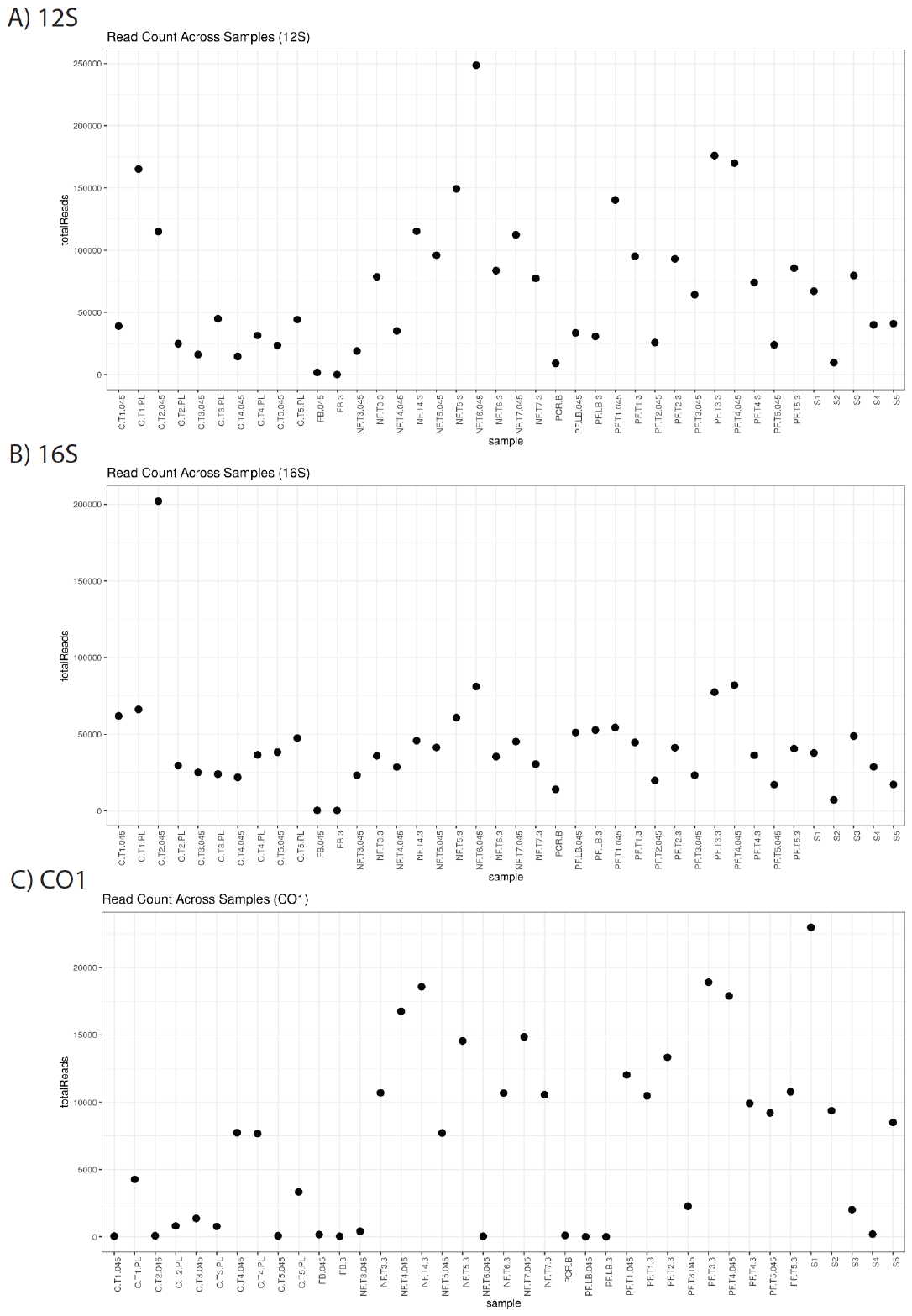

Figure S2: Total number of reads per sample per barcode. C: centrifugation; NF: no freezing; PF: pre-freezing; S: sediment. Numbers at the end of the protocols relate to the size of the filter pores (3 µm and 0.45 µm). PL relates to the extracted pellet from the centrifugation protocol.

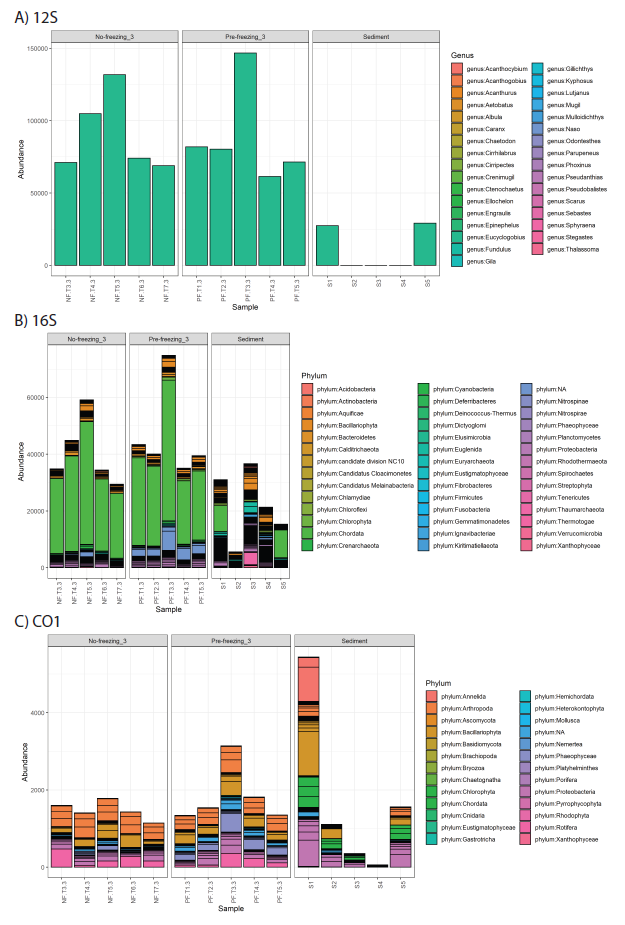

Figure S3: Barplots of genus (A: 12S) and phylum (B: 16S; and C: CO1) read abundance. Sample code the same as Figure S2.

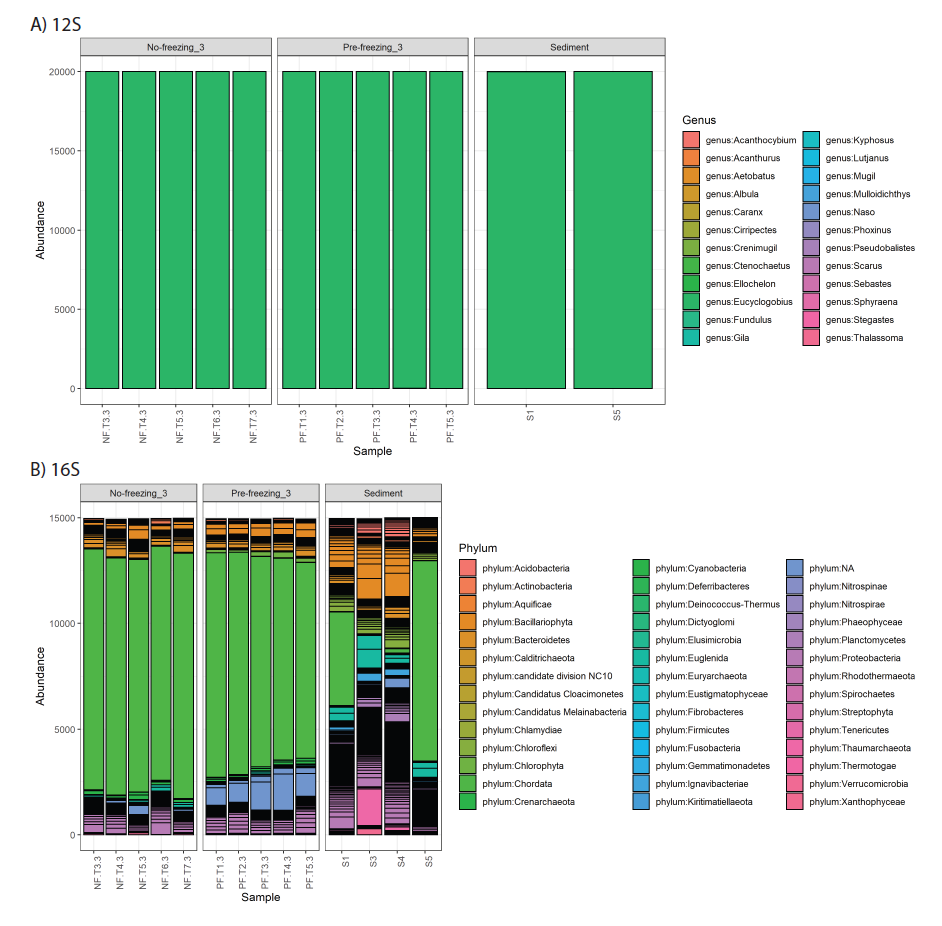

Figure S4: Barplot of rarefied dataset of genus (A: 12S primer) and phylum (B: 16S primer) abundance for each protocol: no freezing, pre-freezing and sediment. Sample code the same as Figure S2.

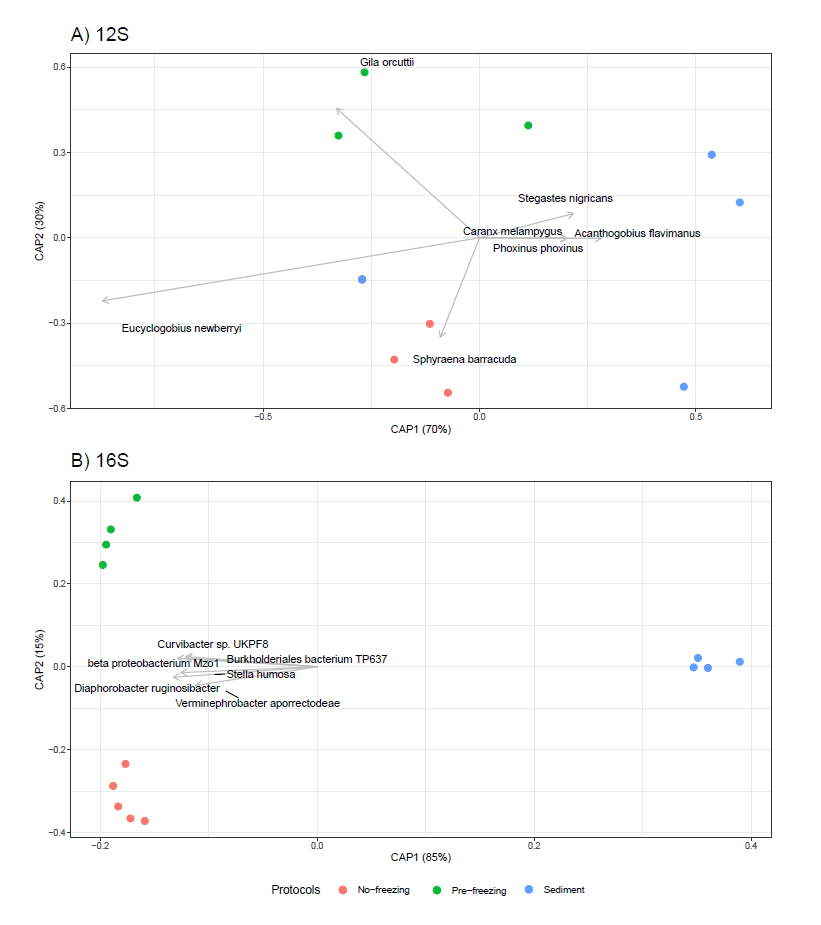

Figure S5: Constrained Analysis of Principal Coordinates (CAP) of A) 12S and B) 16S primer datasets standardized via Wisconsin double standardization (eDNA index). It can be noted that in the 12S dataset, the tropical species suspected to be contaminants from tag-jumping are driving the community assemblage differences between protocols, which is different from the rarefied dataset (Fig. 9).
