## Supplementary figures and images for "Different approaches to processing environmental DNA samples in turbid waters have distinct effects for fish, bacterial and archaea communities"

### Figure S1

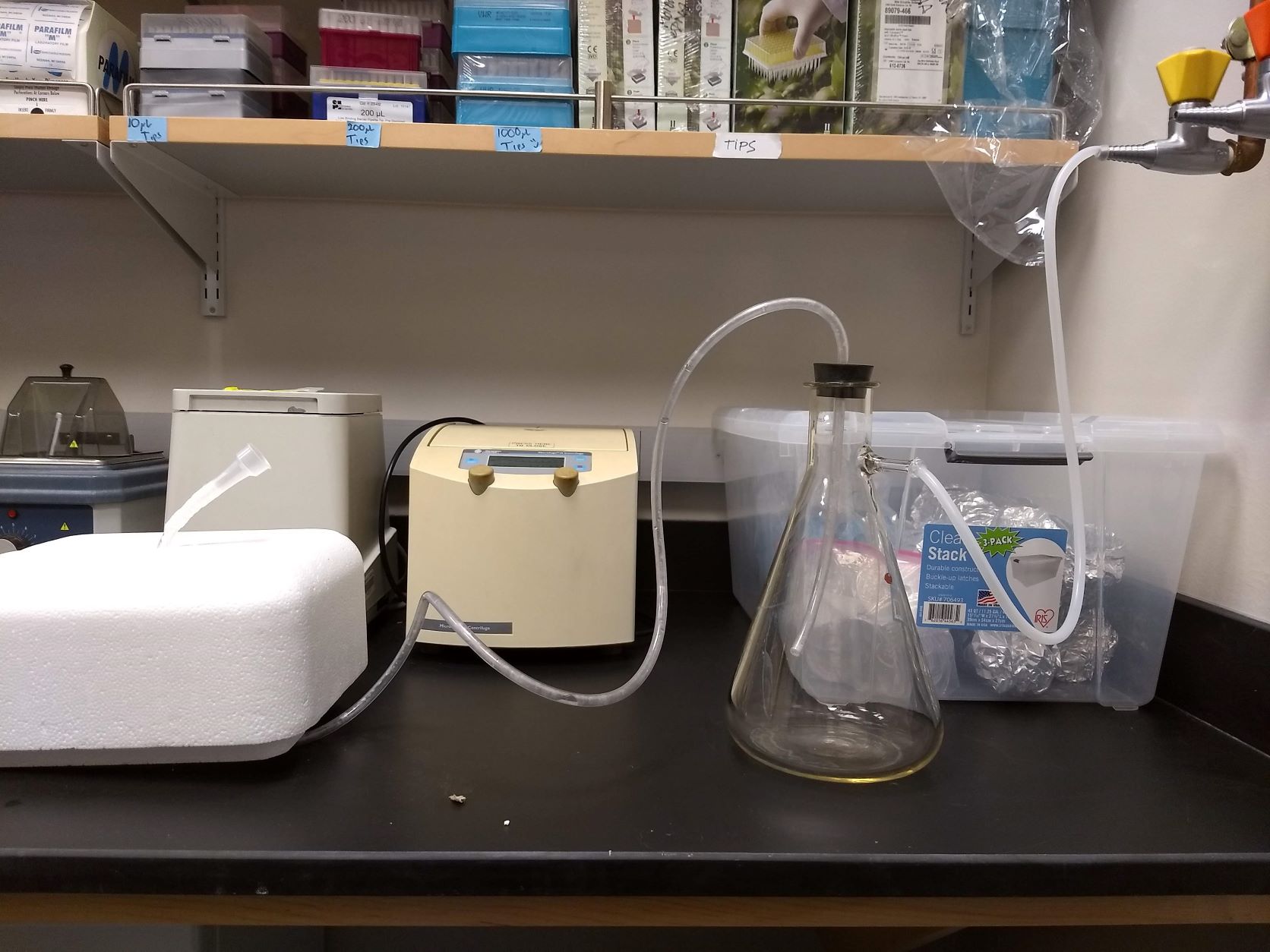

### Figure S2

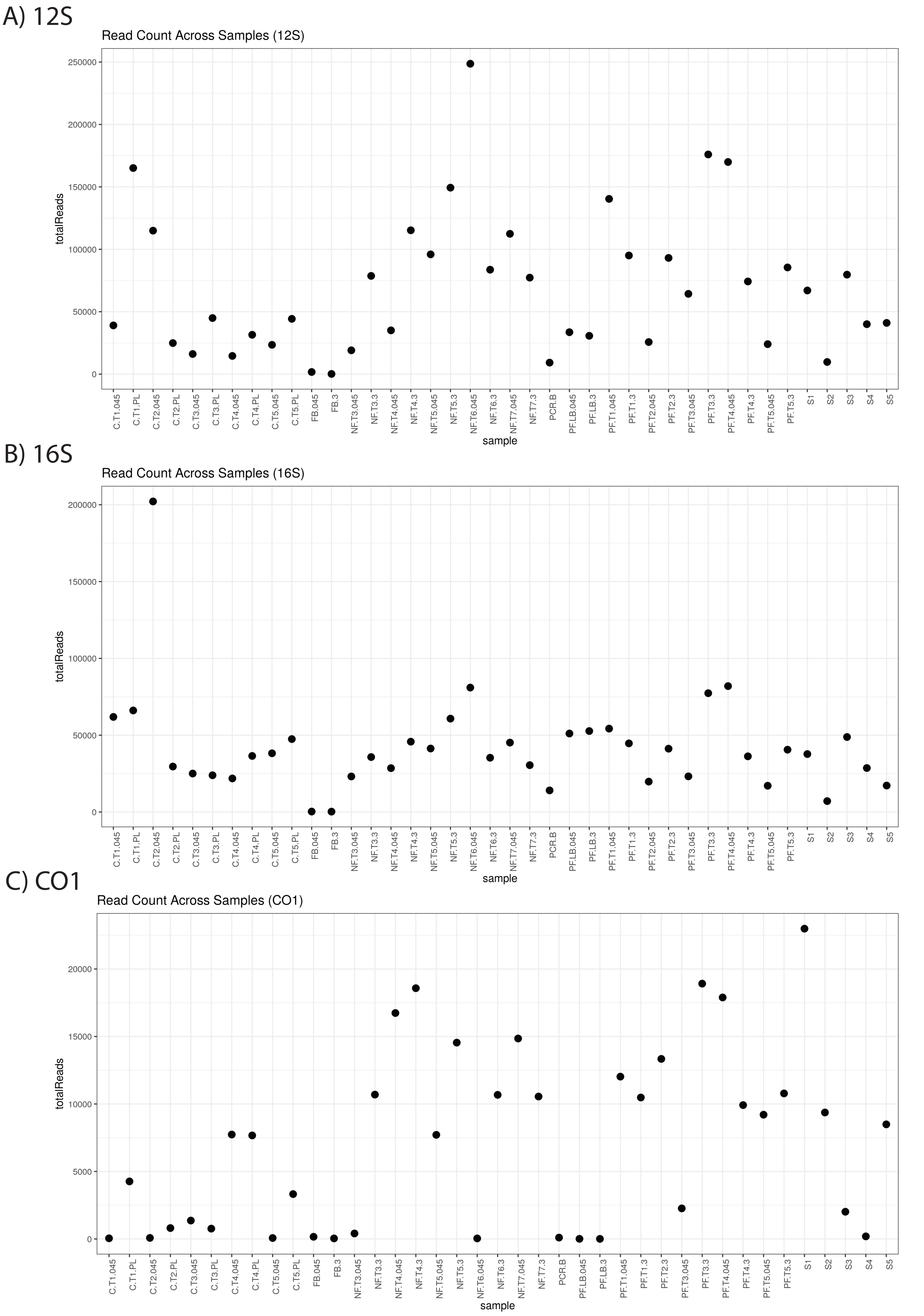

### Figure S3

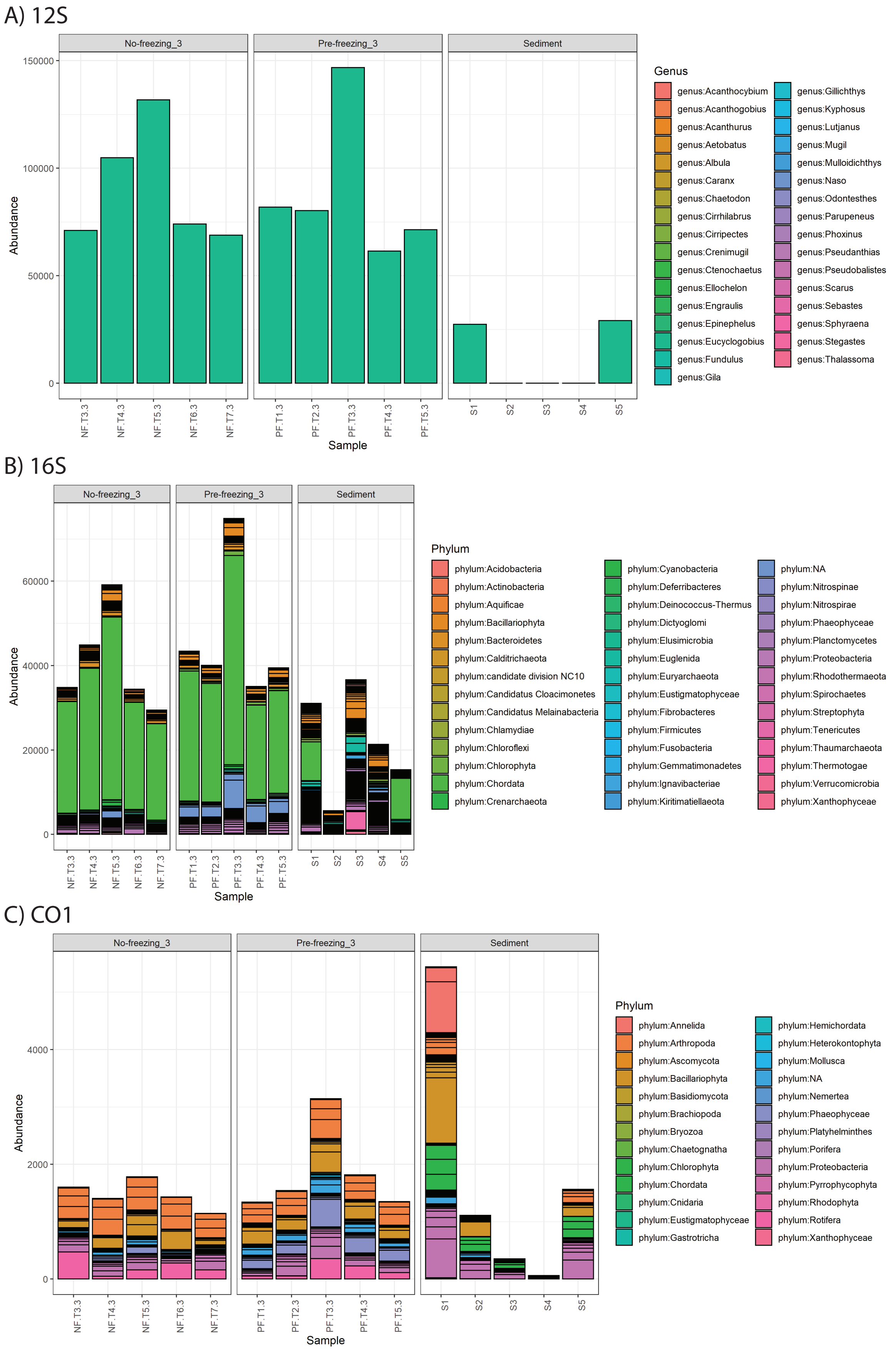

### Figure S4

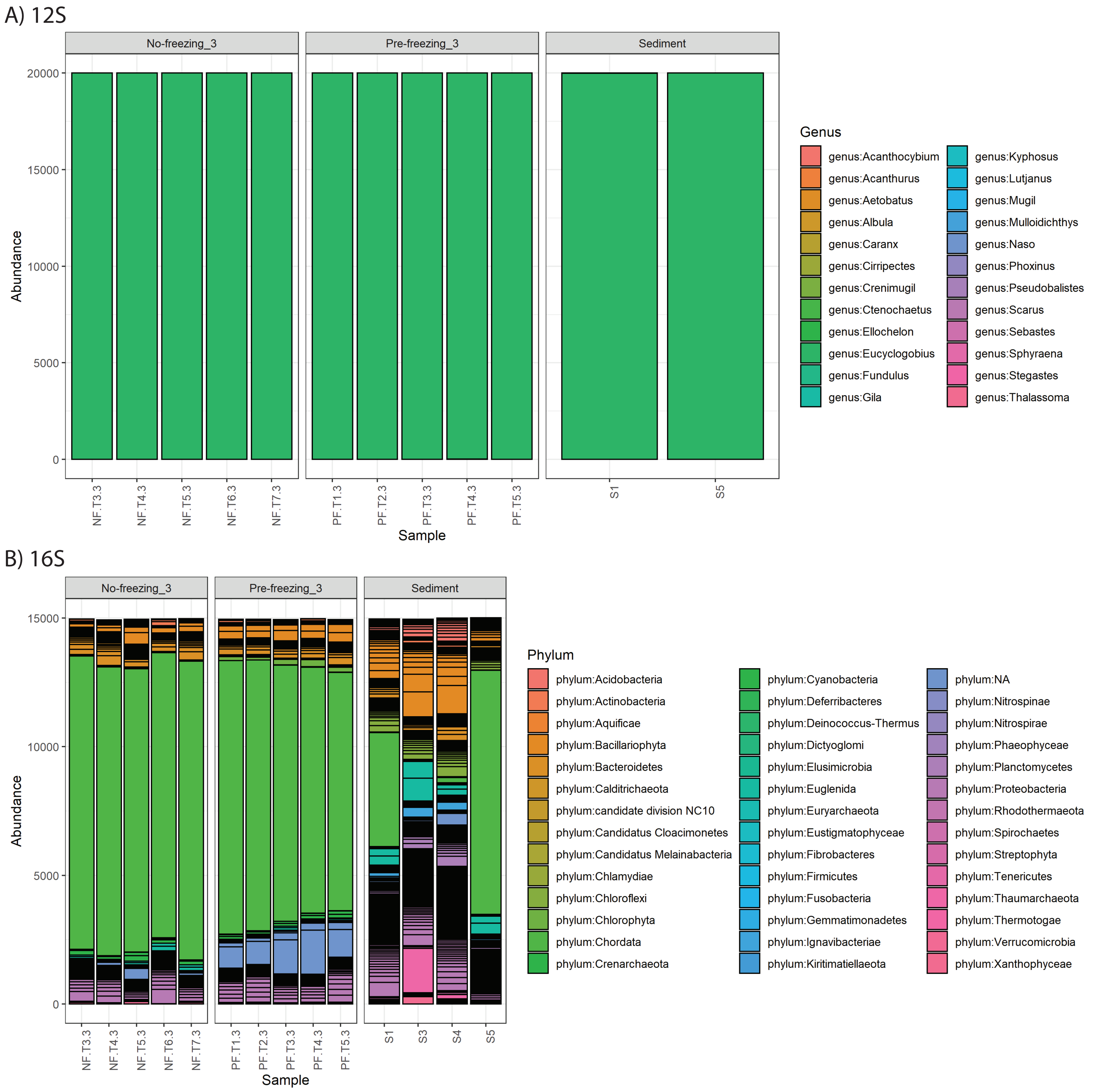

### Figure S5

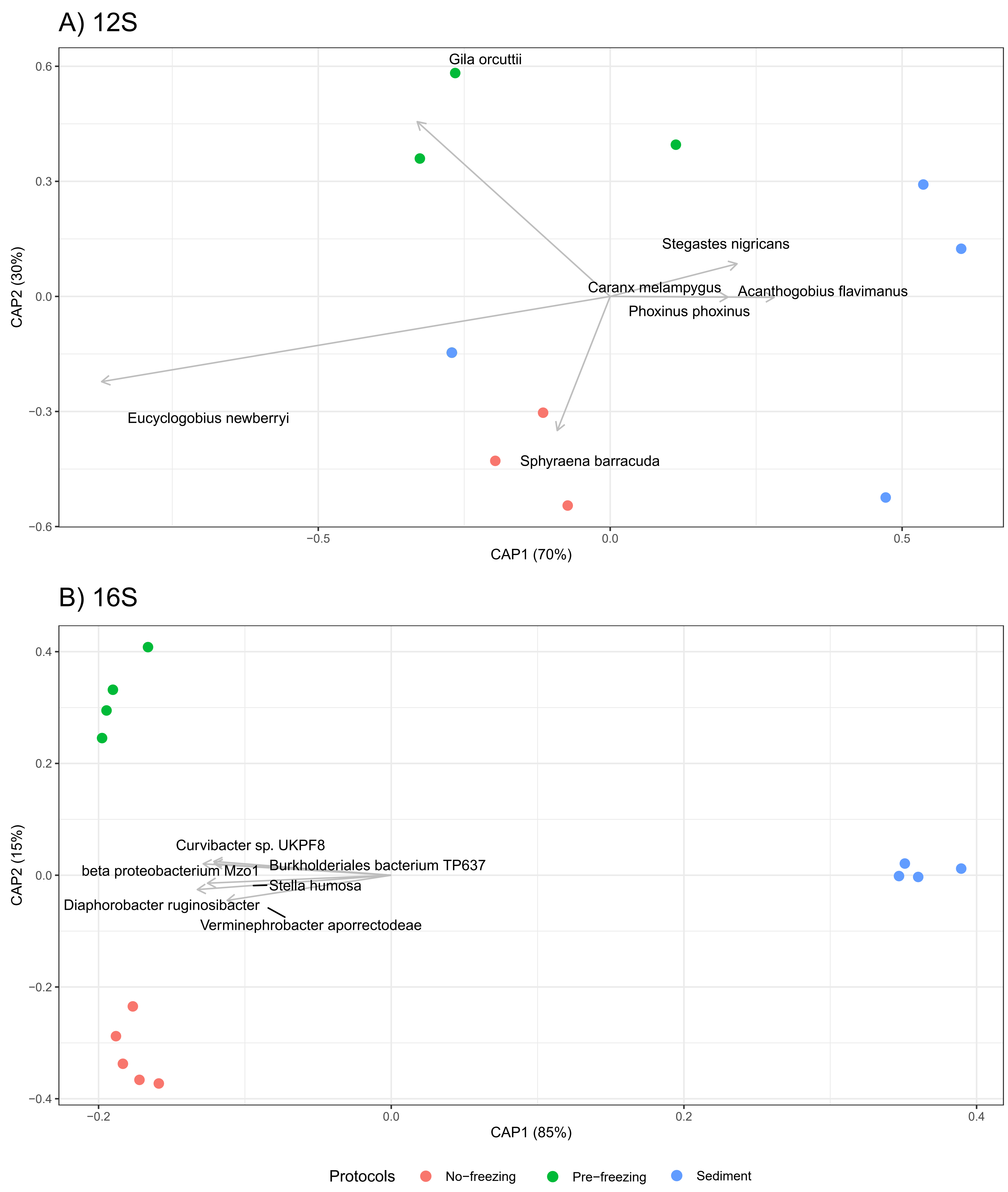
